# Flies adaptively control flight to compensate for added inertia

**DOI:** 10.1101/2022.12.15.520540

**Authors:** Wael Salem, Benjamin Cellini, Eric Jaworski, Jean-Michel Mongeau

## Abstract

Animal locomotion is highly adaptive, displaying a large degree of flexibility, yet how this flexibility arises from the integration of mechanics, sensing and neural control remains elusive. For instance, animals require flexible strategies to maintain performance as changes in mass or inertia impact stability. Compensatory strategies to mechanical loading are especially critical for animals that rely on flight for survival. To shed light on the capacity and flexibility of flight neuromechanics to mechanical loading, we pushed the performance of fruit flies (*Drosophila*) near its limit and implemented a control theoretic framework to quantify how flies compensated for added inertia. Flies with added inertia were placed inside a virtual reality arena which enabled free rotation about the vertical (yaw) axis. Adding inertia increased the fly’s response time yet had little influence on overall gaze performance. Flies maintained stability following the addition of inertia by adaptively modulating both visuomotor gain and damping. In contrast, mathematical modeling predicted a significant decrease in flight stability and performance. Adding inertia altered saccades, however flies compensated for the added inertia by increasing yaw torque production, indicating that flies sense that they are mechanically loaded. Taken together, in response to added inertia flies trade off reaction time to maintain flight performance through adaptive neural modulation. Our work highlights the flexibility and capacity of motor control in flight.

## Introduction

Organisms display a wide array of compensatory strategies to maintain function and performance. Compensatory strategies to mechanical loading are particularly important for flying animals that rely on stable flight for finding food, mate, escape predators, etc. In flying insects, the most drastic weight fluctuations can arise from feeding (Muijres et al., 2017b) and carrying loads (Mountcastle et al., 2015), and can triple overall weight in some cases (van Veen et al., 2020). Previous studies have investigated the robustness of flying insects to small changes in weight or inertia, e.g., (Combes et al., 2020), but the underlying neuromechanical control strategies used to maintain performance are not well understood. Pushing flying insects beyond minor changes in weight, in conjunction with a control theoretic framework, could unravel modes and control strategies that are obscured under natural conditions (Salem et al., 2022). This approach in turn could provide unique insights into the capacity of the nervous system outside the natural context and the role that different sensory modalities play in flight compensation. Indeed, pushing insects beyond their natural context has been fruitful to study the neuromechanics of locomotion on land and in air (Jindrich and Full, 2002; Revzen et al., 2013; Ristroph et al., 2010; Ristroph et al., 2013).

Here, we used system identification to examine the impact of added yaw inertia on the flight performance of tethered fruit flies free to rotate about the yaw axis. To quantify compensatory strategies, we perturbed the gaze stabilization response of flies by placing flies inside a virtual reality flight simulator. Our paradigm allowed flies to close the loop between visual stimulus and their gaze by rotating about the yaw axis. The yaw inertia of fruit flies was altered by mounting 3D printed cylinders with distinct inertia onto the magnetic pin. This paradigm pushed the performance of flies beyond natural conditions as lift generation and yaw stabilization were decoupled, thus providing insights into the capacity of the nervous system to adapt yaw steering. By increasing the yaw inertia of fruit flies by up to sixty-four times (64X), we found that altering inertia had a noticeable impact on both the performance and timing of the yaw gaze stabilization response. Using a control theoretic framework, we demonstrated that adding inertia did not significantly alter the yaw response of fruit flies but intriguingly resulted in a larger response time. Flies maintained similar performance across range of added inertia by increasing both damping and visuomotor gain, likely through the integration of visual and mechanosensory feedback.

## Results

### Flies maintained similar performance at the expense of increased response time to stabilize gaze

Fruit flies were tethered to a magnetic pin and placed inside a virtual reality arena. This configuration restricted the motion of flies to rotation about the yaw axis. The yaw inertia of fruit flies was altered by mounting small 3D-printed cylinders of distinct sizes onto the magnetic pin (Figure 1A–C & Methods). To ensure our study spanned a wide range of added inertias, we designed eight cylinders with logarithmically increasing yaw inertias (see Methods). The smallest cylinder was approximately the same yaw inertia of a fruit fly (5.2×10^-13^ kg m^2^), whereas the yaw inertia of the inertia of the largest cylinder was approximately 64X that of a fruit fly. To measure the impact of altering inertia on flight performance and stability, we first investigated how increasing inertia altered yaw stability in the presence of a static visual panorama. Failure to maintain a stable heading following changes in inertia could indicate a decrease in flight stability. Increasing the yaw inertia by 16X or more caused flies to oscillate about the yaw axis (Figure 1D & Movie S1). With increasing inertia, the magnitude and frequency of these oscillations increased and decreased, respectively (Figure 1E). In contrast, flies with no added inertia did not exhibit such large oscillations (Figure 1D,E). The presence of such oscillations suggests that flies’ stability is impaired by adding inertia beyond a certain amount.

**Figure 1.**
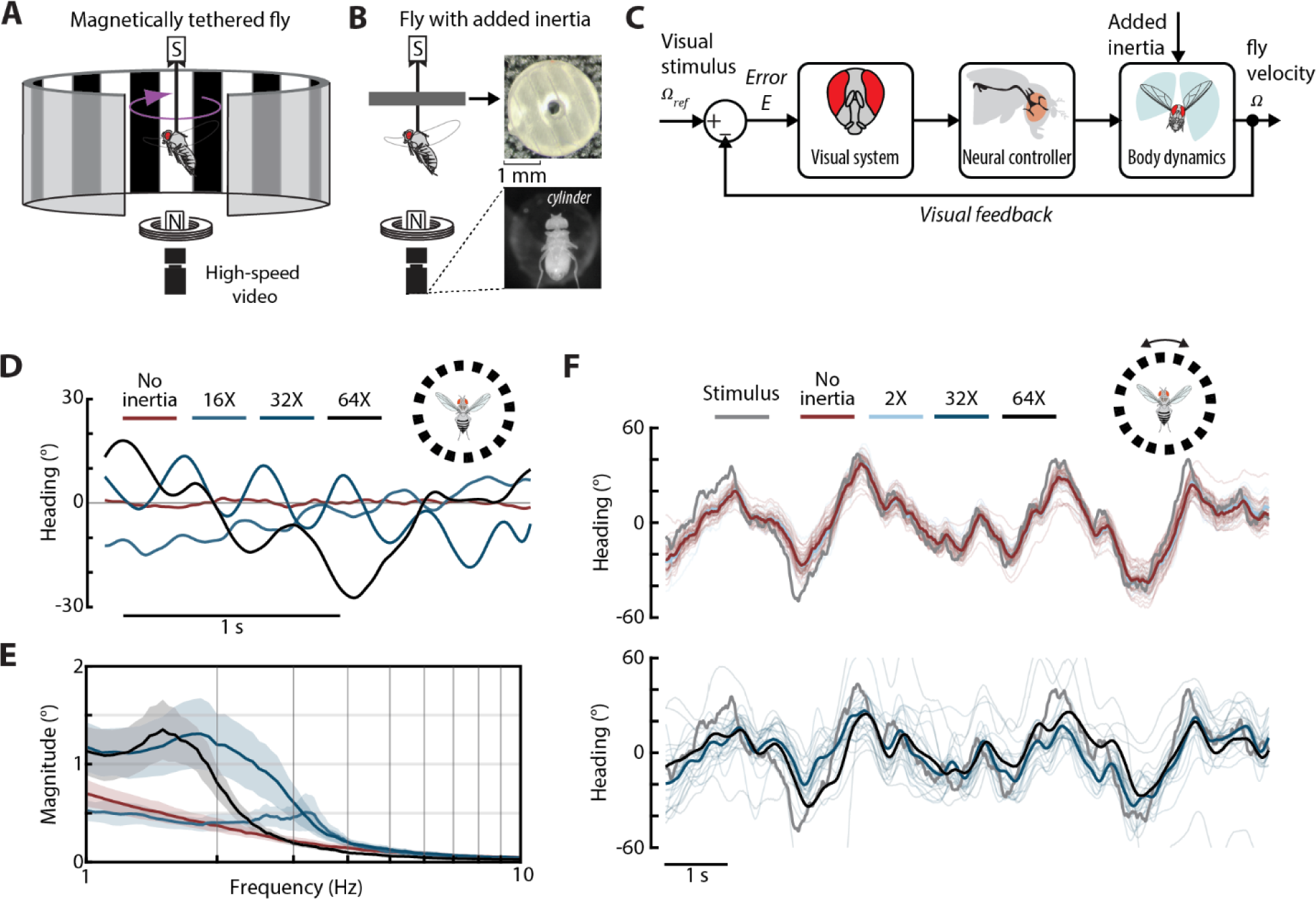
Experimental setup and paradigm to test the impact of increasing yaw inertia on the performance and stability of fly flight. **A)** The magnetic tether system and virtual reality arena. The flies were glued to a magnetic pin and suspended between two magnets inside the virtual reality arena. This configuration enabled rotation about the yaw axis while restricting motion in other directions. Changes in the fly’s heading were recorded using a bottom view high-speed camera. **B)** An illustration of a magnetically tethered fly with a cylinder glued onto the magnetic pin (left). The cylinders (top right) were 3D printed and mounted onto the magnetic pins to increase yaw inertia (bottom right). **C)** The proposed control framework used to model the optomotor response of magnetically tethered flies. **D)** Sample data of individual flies presented a static visual stimulus with different added inertia. **E)** Magnitude plot showing the average frequency and amplitude of the oscillations. **F)** The visual sum-of-sines stimulus (grey) along with the individual fly response (thin lines) and the mean response across all individuals (solid lines) for select amounts of added inertia. For **E**, shaded region is ±1 STD. No added inertia *n* = 13 flies; 16X: *n* = 7 flies; 32X: *n* = 9 flies; 64X: *n* = 13 flies. For **F**, no added inertia: *n* = 41 flies; 2X: *n* = 14 flies; 32X: *n* = 17 flies; 64X: *n* = 8 flies.

To quantify the impact of increasing inertia on flight performance, we presented flies with a sum-of-sines visual stimulus composed of nine sine waves with distinct frequencies and phase (see Methods & Movie S2). The stimulus produced an optomotor response in all the tested groups with and without added inertia (Figure 1F & Figure S1A). Interestingly, even flies with added inertia up to 64X stabilized the moving background. However, a closer inspection of the time domain data revealed a change in performance when the fly’s inertia was increased by 16X or more, consistent with our findings for flies presented a static visual stimulus (Figure 1F & Figure S1). When the inertia of flies was increased by more than 8X, the average response appeared smoother due to the attenuation of the higher frequency components, suggesting that adding inertia primarily influences high-frequency gaze stabilization performance. At the highest tested inertia (64X), the optomotor response was significantly attenuated and no longer coherent with the visual stimulus at most frequencies (Figure Supplement 1B). Therefore, we excluded data collected at this inertia from further frequency domain analysis.

To further examine the impact of added inertia on flight performance, we conducted a frequency domain analysis of the responses to sum-of-sines visual stimuli. By computing gain and phase difference, we mathematically quantified the changes in performance at each frequency component of the stimulus. First, we predicted how the performance of flies might change without changes in controller parameters (neural control). We modeled the yaw body dynamics of the fly as a first-order system (Figure 1C) of the form

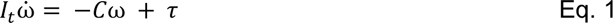

where, *I_t_* is the total yaw inertia of fruit flies (fly inertia + cylinder inertia), *C* is the yaw damping, ω is the yaw angular velocity, and τ is the yaw torque produced by the fly. This modeling assumption is consistent with the notion that fly flight about yaw is damping dominated (Dickson et al., 2010). By assuming that the visual system acts primarily as a proportional gain on velocity—consistent with previous studies that showed little contribution of integral feedback during yaw gaze stabilization maneuvers (Cellini et al., 2022; Salem et al., 2022)—the open-loop transfer function 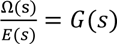 can be written as:

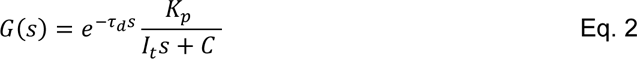

where *s* is the complex frequency, *E*(*s*) is the Laplace transform of the error (velocity) between the stimulus and fly motion, *K_p_*is the visuomotor gain, and τ_*d*_is time delay due to neural processing. Increasing the inertia in Eq. 2 should alter the pole location and consequently the stability of the system. Therefore, increasing yaw inertia without changing damping would push the open-loop pole towards the imaginary axis. When considering the closed-loop system—which is expressed as *G*(*s*)⁄1 + *G*(*s*)—and considering constant proportional gain, increasing inertia increases the system’s time constant (ratio of inertia to damping), and thus a large added inertia could push the system close to a marginally stable or unstable state (Aström and Murray, 2010).

By simulating an increase in inertia (without changing other parameters) and using experimentally determined constants for τ_*d*_, *K_p_*and *C* (Eq. 2)(Salem et al., 2022), we predicted the closed-loop frequency response of flies with added inertia. Importantly, baseline parameters were estimated from flies with no added inertia (See Methods for details). We predicted that the gain would significantly drop at all frequencies if the inertia was altered without changing the baseline damping and visuomotor gain (Figure 2A). We further predicted that the phase difference would decrease significantly at the lower frequencies but converge to the same value at higher frequencies (Figure 2A). The experimentally measured frequency domain response did not resemble our prediction (Figure 2B). The gain was almost unchanged for all inertias up until 0.9 Hz, and the phase difference decreased far beyond our predicted limit (Figure 2B top panel, Table S1, Figure S2). Because mechanics alone could not account for the experimental data, our results strongly imply that flies tune internal gains to compensate for added inertia. At frequencies higher than 0.9 Hz, the difference in gain among the inertia altered flies became statistically significant (Table S1) but did not follow a specific pattern. In fact, flies that had their inertia increased by 16 and 32 times had the largest gains at 1.45 Hz and 2.25 Hz. However, the increase in inertia began to reduce the gain at frequencies above 3.45 Hz. At frequencies higher than 3.45 Hz, the gain was significantly smaller at all added inertias compared to the intact case (Table S1). Interestingly, the response when the yaw inertia was increased by 32X resembled the response of a second-order underdamped system rather that of a first-order system. The gain peaked at 2.25 Hz and was greater than unity. However, the frequency at which the peak occurs coincides with the frequency of observed oscillations with a static stimulus (Figure 1E, Figure 2B). This peak complicated the interpretation of the gain data at this inertia as it could be a result of superimposed noise or due to higher order dynamics.

**Figure 2.**
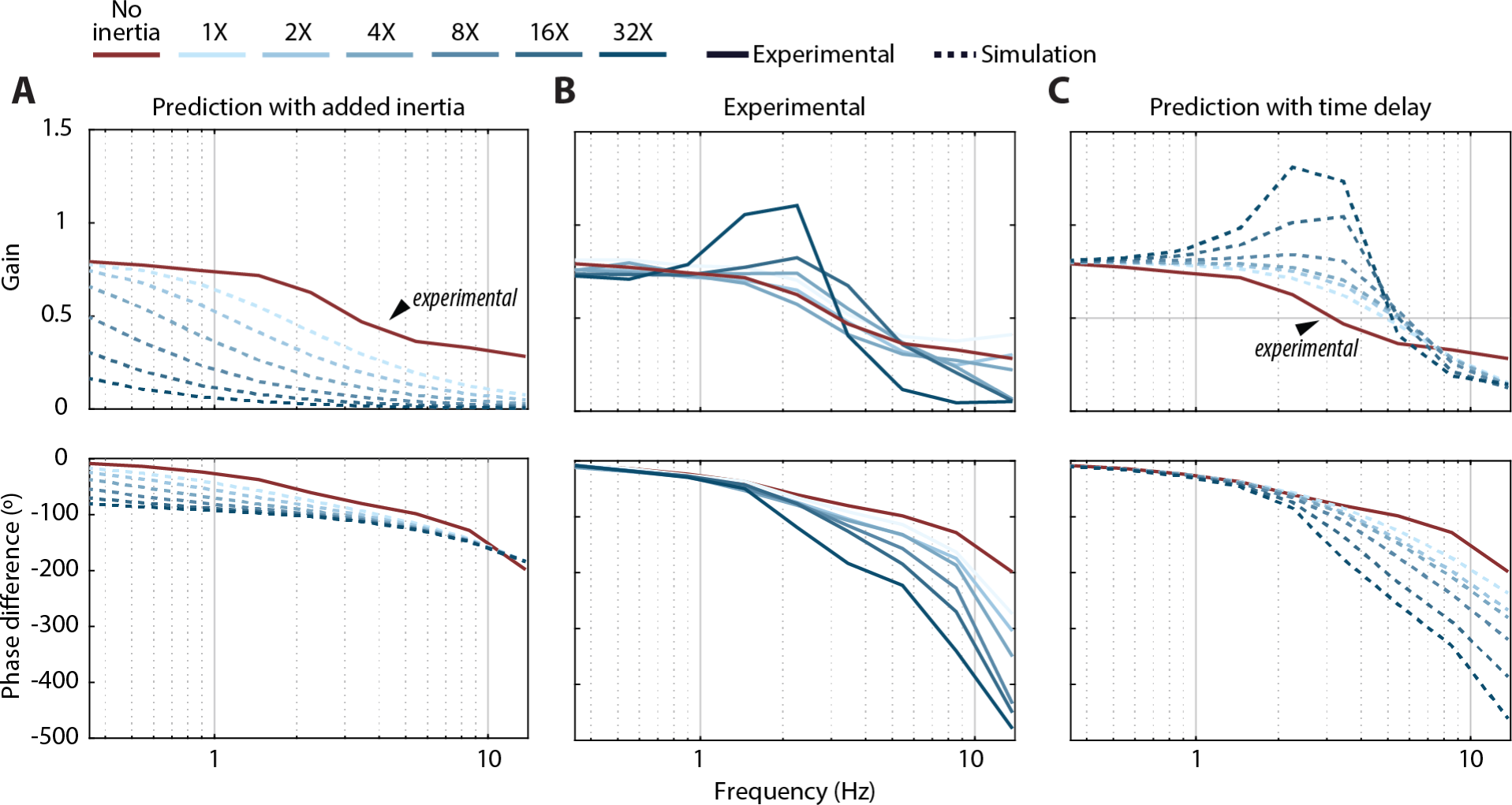
Flies maintained similar performance at the expense of increased response time to stabilize gaze. **A)** The average experimental closed-loop response with no added inertia (red line) versus the simulated response to additional yaw inertia (dashed lines). **B)** The empirical frequency response function of flies with added inertia in response to a sum-of-sines stimulus. Addition of inertia had a significant influence on the phase difference and gain for frequencies greater than ∼0.9 Hz (see Table S1 for exact values and statistics). **C)** Simulated frequency response functions for a no-added-inertia fly with increasing time delay. For **A and C,** dashed lines are from simulation and solid lines are the experimental results. Plots with ±1 STD are shown in Figure S2. No inertia added: *n* = 41 flies; 1X: *n* = 11 flies; 2X: *n* = 15 flies; 4X: *n* = 19 flies; 8X: *n* = 17 flies; 16X: *n* = 17 flies; 32X: *n* = 8 flies.

Looking at the phase data can shed light on the underlying cause behind this rise in gain. Increasing inertia altered the phase difference in a completely different manner than predicted by simulation (Figure 2A,B). The variation in phase difference became more significant at larger inertias for stimulus frequencies at and above 1.45 Hz (Figure 2B lower panel; Table S2). Such changes in phase cannot be explained purely by altering the damping or inertia in our model (Eq. 2). In the absence of a time delay, the phase difference of first-order systems converges to −90° at high frequencies. Hence, the observed changes in phase difference are likely a result of an increase in the time delay (τ_*d*_). Simulating Eq. 2 with no added inertia and delays estimated from the empirical phase difference (see Methods) captured the changes in both the gain and phase difference (Figure 2C). Altering the time delay even captured the peak observed at 32X added inertia. The interpretation of this result is not intuitive as time delays are usually associated with changes in phase difference but not gain. However, altering the delay in the open-loop system influenced both the gain and phase of the closed-loop system. Changes in the time delay of the intact system could capture our empirical results, however this simulation did not account for changes in inertia and did not consider how the remaining internal parameters are modulated to maintain the same gain. Altogether, our findings demonstrate that flies can compensate for an increase in inertia, with the trade-off of increased time delay.

### The head compensates for loss of stability but not for changes in gaze performance

We demonstrated that increasing the yaw inertia of flies by more than eight times caused a modest change in flight performance and stability. When presented with a visual sum-of-sines stimulus, fruit flies experienced a small decrease in gain and phase at frequencies larger than 1.45 Hz. On the other hand, flies presented with a static stimulus began to oscillate about the yaw axis and failed to maintain stable body heading. Taken together, these results indicate that the body response of flies significantly deteriorated with the addition of inertia. However, this may not be the case for the overall gaze as head motion could be used to compensate for changes in body motion (Cellini and Mongeau, 2020a). Previous work explored the overall role of the head in gaze stabilization (Cellini and Mongeau, 2020a). The head compensates for fast visual motion, whereas the body compensates for slower visual motion. By combining head and body motion, flies can improve overall gaze stabilization performance over a large range of visual motion velocity (Cellini et al., 2022). Thus, the head may adjust its motion to compensate for changes in stabilization performance following the addition of inertia to the body.

To gauge the compensatory role of the head, we tracked the motion of the head in experiments with a sum-of-sines and static background stimulus. We then conducted a frequency domain analysis to determine the response of the head (Figure S5A). When presented with a sum-of-sines stimulus, the head did not appear to play a large compensatory role. The gain and phase difference of the head were similar across all groups of flies. While the phase difference at the highest three frequencies fluctuated among different inertia treatments, it did not follow a clear trend. This is likely due to the fact that body motion influences the visual input to the head controller, and body motion varies greatly at high frequencies for different added inertia (Figure 2B). Thus, changes in visual feedback likely led to these changes in head phase. Of interest was the peak in gain observed at ∼2.3 Hz (Figure S5A), which coincided with the peak observed in body gain at the same frequency (Figure 2B). Changes in head gain at this frequency are likely an attempt to compensate for elevated oscillatory body motion within this range of frequencies.

In stark contrast, flies presented with a static stimulus altered head motion to compensate for body instabilities. We previously showed that flies oscillated about the yaw axis when their inertia was increased by 16X or more, and the frequency of these oscillations were dependent on the amount of added inertia (Figure 1D,E). To compensate for these body oscillations and maintain stable gaze, flies increased head motion at those frequencies, presumably to cancel out body motion (Figure S5B). However, the change in head motion was not sufficient to completely cancel body motion (Figure S5B). Interestingly, the time difference between the head and body increased with larger inertias (Figure S5C) (*p*\*\*\*, ANOVA; DoF = 3).

### Flies increased visuomotor gain and damping to maintain the same stability

We demonstrated that increasing inertia noticeably altered the performance and timing of the optomotor response especially at higher frequencies (Figure 2B). From a control perspective, increasing the inertia of a first-order system with a proportional controller shifts the pole of this system closer to the imaginary axis, and causes a significant drop in the gain and phase difference (Figure 2A). A closed-loop system could possibly compensate for changes in stability by altering its controller through an adaptive control scheme. However, if the goal is to maintain the same performance and stability, a change in proportional control alone cannot produce the desired change in the system’s dynamics.

To shed light on how increasing yaw inertia altered the yaw dynamics of flies, we fit the empirical frequency response functions (FRFs) (Figure 2B) to a first-order transfer function with a delay, one pole, and no zeros (Cellini et al., 2022). Specifically, we used a least square estimate to fit the open-loop transfer function *G*(s) of flies with and without added inertia (Figure S3A) (Roth et al., 2012), where the open-loop transfer function is of the form shown in Eq. 2. There was some individual variation between animals when fitting, and this variation became more prominent at the higher inertias. However, a first order model captured the open-loop dynamics of the optomotor response of all groups (r-squared ∼88%, see Figure S3B). To verify that our fit model properly captured the time domain response, we simulated the fit transfer functions using the same sum-of-sines visual stimulus as the input. The simulated response closely resembled the actual response of flies (Figure S4).

Estimating the open-loop parameters can shed light on the underlying neuromechanical control strategies used by flies to compensate for changes in inertia. The numerator is the visuomotor gain which could be modulated by the fly. On the other hand, the denominator is used to determine the location of the open-loop pole and thereby measure the stability of the open-loop system. Finally, the delay term can provide an estimate of the system’s lag due to sensorimotor processing. Comparing the various fits, we first noticed that changes in the system time delay were positively correlated to changes in inertia (Figure 3A). With no added inertia, the time delay was around 20 ms, which is consistent with previous studies (Cellini et al., 2022). However, this delay steadily increased with increasing inertia, up to approximately 80 ms (Figure 3A).

**Figure 3.**
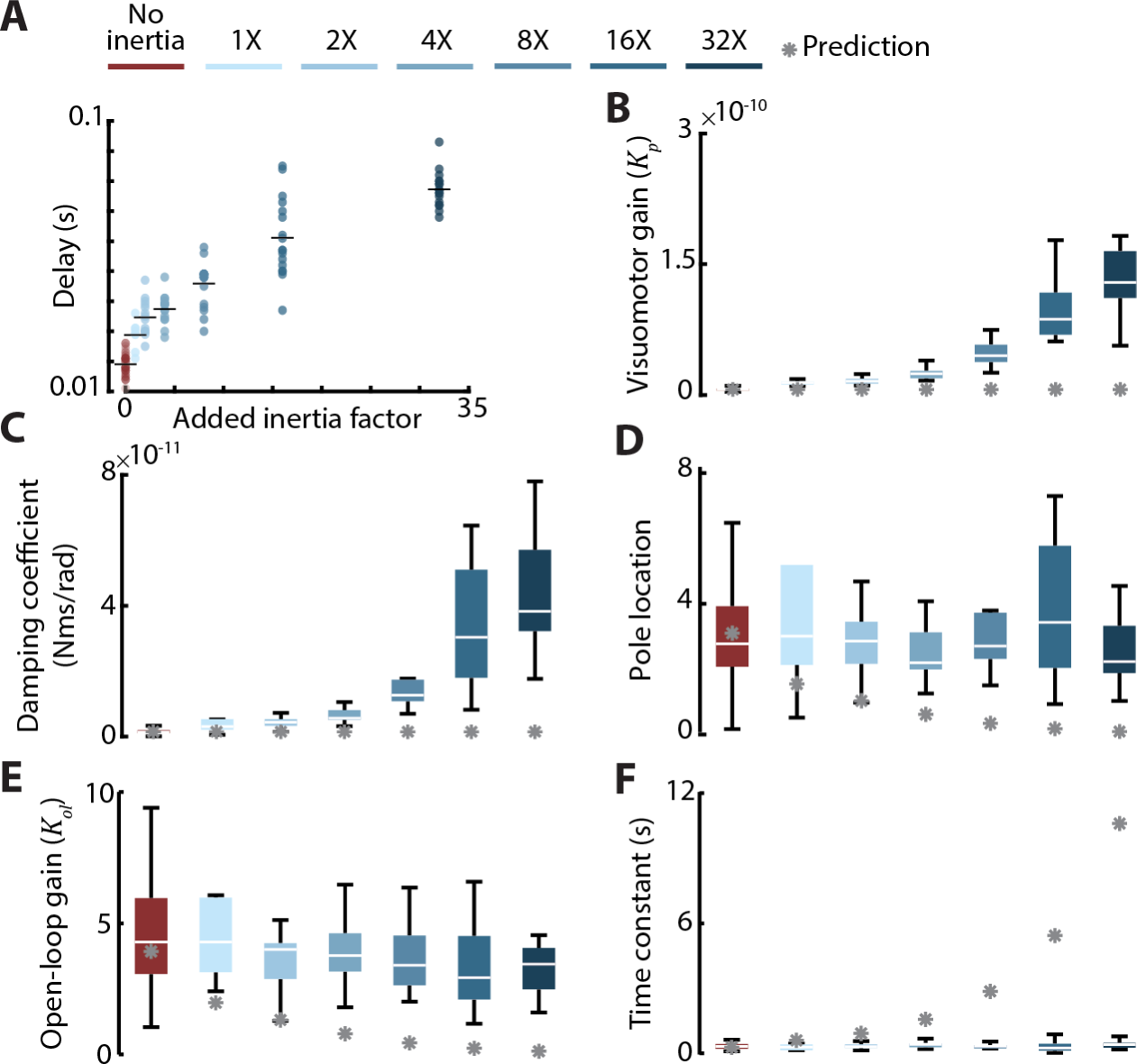
Flies increased visuomotor gain and yaw damping to maintain stability. **A)** Estimate of the time delay for intact flies and flies with added inertia. Increasing inertia caused the time delay to increase. This increase was proportional to the amount of added inertia (ANOVA; DOF = 6; *p* < 0.001). Horizontal line: average. **B)** The predicted (grey asterisk) and estimated visuomotor gain. In response to increases in inertia, flies modulated their visuomotor gain (ANOVA, DOF = 6; *p* < 0.001). **C)** The estimated active damping. Flies actively modulated their yaw damping (ANOVA; DOF=6; *p*<0.001). **D)** Pole location of flies with different added inertia. The overall pole location of fruit flies changed marginally (ANOVA; DOF = 6; *p* = 0.04). **E)** The open-loop gain (*K*_*ol*_) and **F)** the system time constant (τ_*f*_). Flies maintained the same open-loop gain and time constants. ANOVA, DOF = 6, *p* = 0.53 & *p* = 0.94, respectively. For all panels: No inertia added: *n* = 41 flies; 1X: *n* = 11 flies; 2X: *n* = 15 flies; 4X: *n* = 19 flies; 8X: *n* = 17 flies; 16X: *n* = 17 flies; 32X: *n* = 8 flies. Further details on goodness of fit and pole locations can be found in Figure S4. For **B–F:** Grey asterisks are the prediction from unaltered fly model parameters (Eq. 3) with inertia as the only parameter change.

Concomitantly the visuomotor gain and damping coefficient significantly changed with increasing inertia (Figure 3B,C). The mean visuomotor gain increased by two orders of magnitude from no added inertia to 32X (Figure 3B). Similarly, the damping increased by more than one order of magnitude (Figure 3C). The location of the open-loop pole changed modestly when the inertia of flies was increased by 32X (Figure 3D, Figure S3C). While adding inertia did alter pole locations (*p*=0.04, ANOVA, 6 DoF), the statistical analysis yielded a *p*-value that is marginally significant. Consequently, this statistical significance may be a result of the fluctuations in pole locations at different inertias. Flies drastically modulated their visuomotor gain and yaw damping in response to changes in inertia, however it is difficult to compare overall changes in system dynamics. To get an idea of how the dynamics of the system changed, we divided the fit transfer functions by the damping to obtain the standard form of a first-order transfer function. The open-loop transfer function becomes:

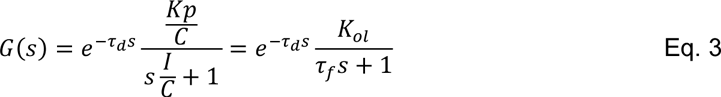

where *K*_*ol*_is defined as the open-loop visuomotor gain, and τ_*f*_is the system time constant. Indeed, estimating the open-loop gain and time constant shows that these two parameters only marginally change (Figure 3E,F). While subject to fluctuations with different amounts of added inertia, the open-loop gain and the time constant remained approximately the same regardless of how much inertia was added (*p*=0.5 & 0.9 respectively; ANOVA, DoF = 6). Thus, flies increased their damping and visuomotor gain to maintain approximately the same open-loop dynamics. Compared to our simulated Bode plots (Figure 2A), flies only experienced a marginal drop in performance following addition of inertia. This suggests that flies modulate both parameters to maintain the same open-loop dynamics. Indeed, by estimating the yaw damping coefficient, we found that flies significantly increased this term to maintain the same pole location, and hence, the same body dynamics. Considering movement of the head—which play a critical role in shaping visual inputs (Cellini and Mongeau, 2020a; Cellini et al., 2021; Cellini et al., 2022)—did not change these conclusions (Figure S5). Therefore, flies maintained roughly the same body dynamics at the expense of a delayed response to the visual stimulus.

### Changes in yaw damping cannot be explained by passive aerodynamics and require active feedback

Using system identification techniques, we found that flies regulated yaw damping to maintain the same open-loop body dynamics following changes in inertia. Damping could be actively modulated though neural control using an inner mechanosensory feedback loop (Figure 4A) (Elzinga et al., 2012; Fuller et al., 2014). Alternatively, changes in damping can be passively regulated as a byproduct of changes in wing kinematics to meet the larger torque requirements imposed by adding inertia. Flapping flight exhibits passive damping about the yaw axis which is generated as a byproduct of drag on wings during flapping. Hence, when flying animals with flapping wings rotate about yaw, a torque is passively produced in the opposite direction of motion. This torque is dubbed flapping counter-torque and damps out turns (Cheng et al., 2010), and can be estimated from wing morphology and kinematics to yield a yaw damping coefficient (Hedrick et al., 2009).

**Figure 4.**
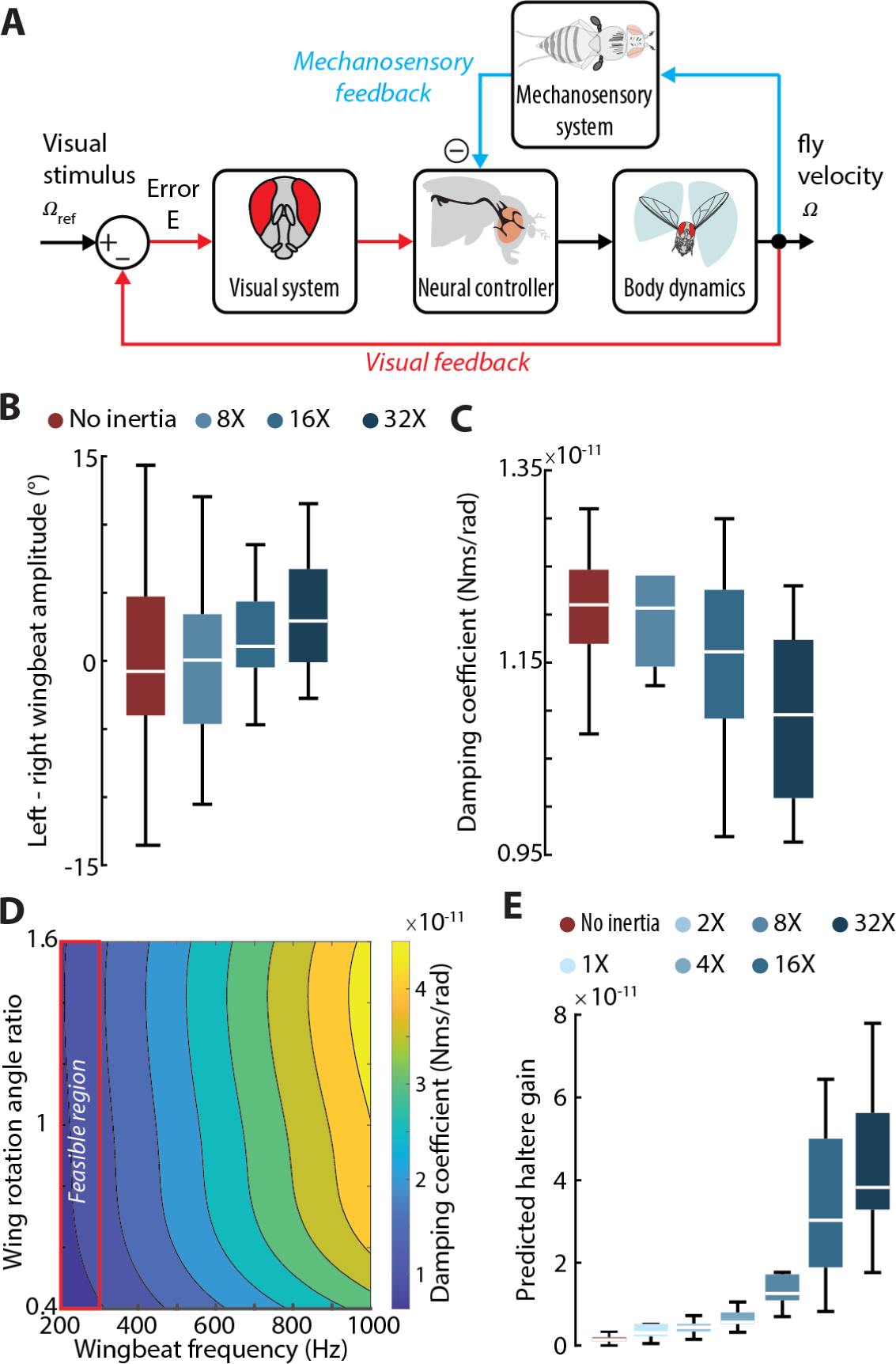
Proposed mechanism for regulating yaw damping. **A)** The proposed control architecture to modulate damping and maintain the same stability. Using the nested mechanosensory feedback loop, flies could alter yaw damping by regulating the gain from the haltere feedback. **B)** The impact of increasing inertia on the difference in wingbeat amplitude (DWBA). Changes in inertia caused no significant changes in DWBA (ANOVA, DOF = 3, *p* = 0.28). **C)** Estimates of the passive damping using a model flapping counter-torque. The model predicted marginal changes in the FCT damping coefficient based on the 2D wing kinematics. **D)** Changes in the FCT as a function of changes in the flapping frequency and magnitude of the wing rotation angle ratio. The magnitude of the rotation angle was modified by multiplying the intact-wing rotation angle with a scaling factor, whereas the frequency was varied from 200 Hz to a 1000 Hz. Red rectangle: region in which flies can feasibly modulate flapping frequency. **E)** Estimated changes in haltere feedback gain in response to changes in inertia. For **B,C, & E:** No inertia added: *n* = 41 flies; 1X: *n* = 11 flies; 2X: *n* = 15 flies; 4X: *n* = 19 flies; 8X: *n* = 17 flies; 16X: *n* = 17 flies; 32X: *n* = 8 flies.

To determine if the change in yaw damping was actively modulated or a by-product of the increase in FCT due to changes in wing kinematics, we first estimated the 3D wing kinematics of magnetically tethered fruit flies with distinct added inertia (see Methods). By Incorporating wing morphology information with wing kinematics of flies with different added inertia (Figure 4B), we found that the FCT does not noticeably change following the addition of inertia if wing beat frequency and rotation angle was to remain the same (Figure 4C). This result may be skewed as flies can regulate wing beat frequency and the rotation angle, hence an increase in modulating these two parameters can significantly influence the FCT. To tease out their relative contribution to the changes in passive yaw damping, we simulated the FCT model (see Methods) and found that flies would need to flap at around 800 Hz to achieve the damping estimated in the FRFs regardless of changes in rotation angle (Figure 4D). This value is much larger than what has been previously reported (Tammero and Dickinson, 2002) and far beyond the physical limit of fruit flies. Therefore, an increase in FCT is not enough to explain the increase in the yaw damping and requires active control.

Fruit flies combine sensory information from multiple modalities to control and regulate flight (Fuller et al., 2014; Sherman and Dickinson, 2004). Therefore, the damping coefficient of the body is likely regulated through an inner sensory feedback loop other than vision (Figure 4A). By integrating an inner loop within the open-loop transfer function *G*(*s*), Eq. 2 can be written as:

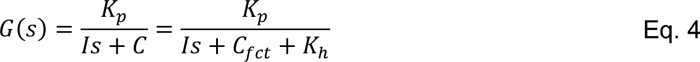

where *C*_*f ct*_ is the passive yaw damping due to flapping counter-torque, and *K*_ℎ_ is the inner loop feedback gain that modulates damping (here we omit the delay term *e*^−τ*d*^_*s*_ for clarity) (Elzinga et al., 2012). The visuomotor gain, *K*_*p*_, can be factored out of Eq. 4 to obtain the formulation of the second inner loop:

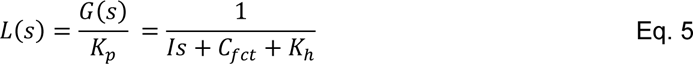

Since the halteres play an important role in encoding angular velocity about the yaw axis and act

faster than vision (Dickinson, 1999), the inner feedback loop is likely driven using mechanosensory feedback from the halteres. Therefore, by modulating *K*_ℎ_ flies could actively increase damping about the yaw axis to maintain the same body dynamics. Changes in the haltere feedback can be estimated by subtracting the FCT from the active yaw damping:

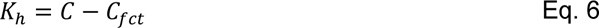

However, this results in negative values of *K*_ℎ_ at 1X inertia which implies that haltere feedback transitions from positive to negative feedback as inertia is increased. To facilitate comparison across all inertia, we shifted our data so that the lowest estimated value of haltere gain was zero. Using this posited control architecture, we estimated that the haltere gain increased with increased inertia to maintain the same yaw damping (Figure 4E). On the other hand, the open-loop visuomotor gain *K_p_*is regulated using visuomotor feedback and maintains the same open-loop performance which would have significantly deteriorated due to elevated inertia and damping. To summarize, our simulation suggests that flies rely on feedback from multiple sensory modalities to maintain the same body dynamics in response to changes in inertia. Our findings hint that flies implement an adaptive control scheme to compensate for changes in inertia. Here, the gains are modulated to maintain gaze stabilization performance.

### Flies compensate for added inertia to control saccades

Saccades are ballistic movements in which flies change their heading in the span of 50–100 ms (Muijres et al., 2015). Such maneuvers have been observed in free and tethered flight (Bender and Dickinson, 2006a; Cellini and Mongeau, 2020b; Land and Collett, 1974). In the magnetic tether, these saccades can be externally triggered from visual cues or internally triggered (spontaneous saccades) (Censi et al., 2013; Mongeau and Frye, 2017). By presenting inertia altered flies with a static stimulus, we measured the impact of inertia on the dynamics of spontaneous saccades. Our simulation (see Methods) predicted that without active control of yaw torque, the displacement, peak velocity, and duration of saccades should greatly diminish following any increase in inertia (Figure 5A bottom panel). Compared to unaltered flies, flies with added inertia exhibited a clear change in saccade dynamics that did not match our prediction (Figure 5A top panel, Figure S6). This was also accompanied by an increase in peak yaw torque during a saccade (Figure 5B). Flies with added inertia exhibited an increase in saccade displacement compared to unaltered flies. This difference became more prominent as the inertia increased (Figure 5C). In contrast, the peak velocity of saccades marginally decreased with increasing inertias (Figure 5D). However, saccade durations exhibited the most change with increasing added inertia (Figure 5D). At no added inertia, the mean saccade duration was just below 100 ms. This value steadily increased with added inertia, and was close to 400 ms when the inertia of flies was increased by 64 times. Complicating this analysis is the large number of samples collected for each inertia treatment, thus tiny differences in saccade dynamics could potentially result in a small *p* value using conventional statistical methods. To address this limitation, we computed Hedge’s g, which is a metric that is independent of sample size (Kelley and Preacher, 2012). Using this effect size model, we found that adding inertia had the largest overall impact on saccade duration and resulted in the largest values of Hedge’s g. As expected, such changes in saccade dynamics required overall higher torques exerted over a longer duration (Figure 5B). Together, these results suggest that flies adaptively control saccade dynamics to compensate for added inertia.

**Figure 5.**
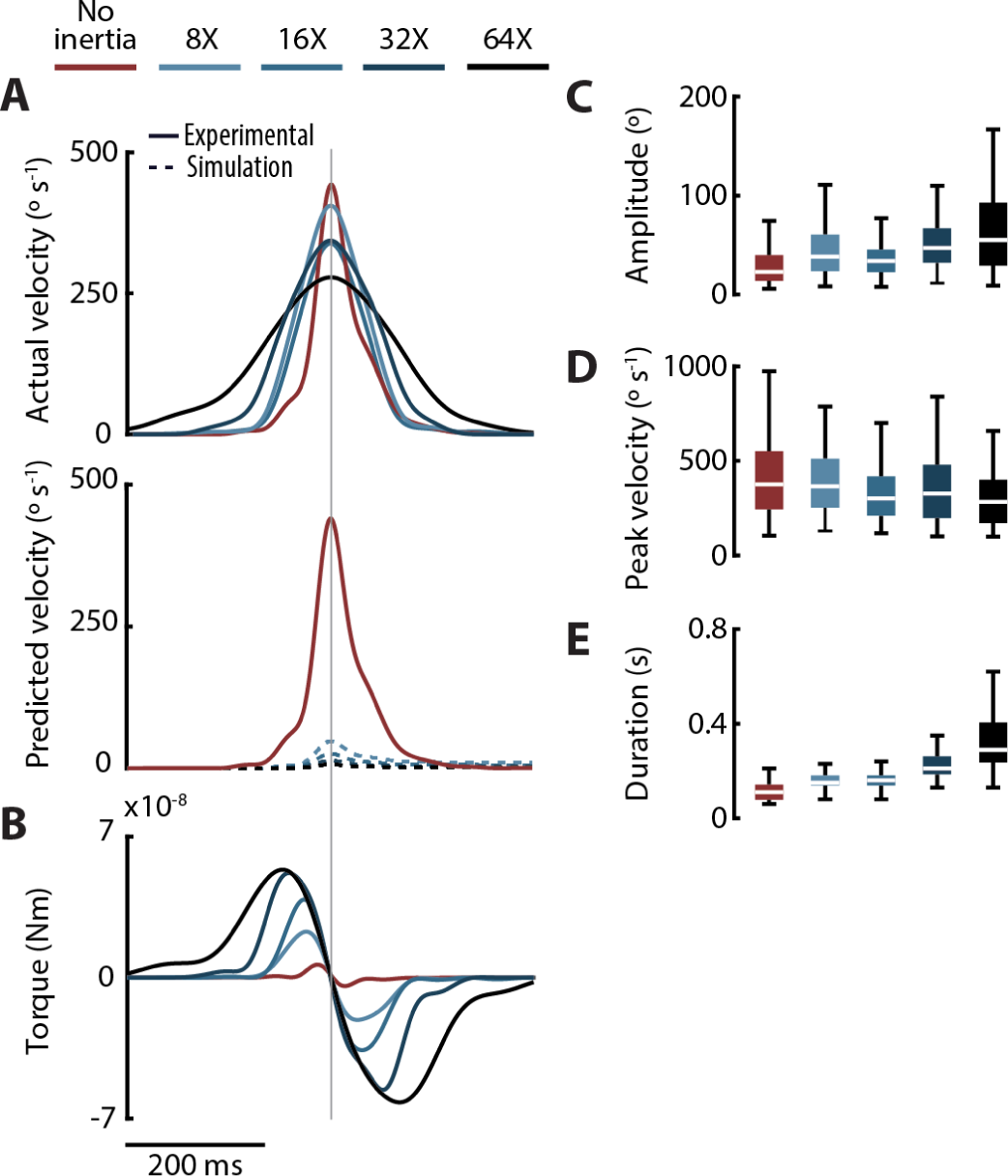
Flies adaptively control saccades. **A)** The average velocity profile of saccades for flies with no or added inertia (top panel), and the predicted saccade velocity profiles estimated using simulation (bottom panel, dashed lines). **B)** Torque profile of flies with added inertia. As more inertia is added, flies produce more torque over a larger duration of time. **C)** Saccade displacement, **D)** peak velocity, **E)** and duration for flies. Adding inertia led to a slight increase in saccade displacement, a slight decrease in peak velocity, and a noticeable increase in saccade duration (Table S4). For **A,B**: Grey vertical line: peak velocity. **For all panels**: No added inertia: *n* = 301 saccades from 13 flies; 8X: *n* = 148 saccades from 9 flies; 16X: *n* = 139 saccades from 7 flies; 32X: *n* = 89 saccades from 9 flies; 64X: *n* = 118 saccades from 13 flies. For saccade variation data see Figure S6.

## Discussion

We discovered that fruit flies adaptively control flight following a large increase in yaw inertia. Specifically, by modulating visuomotor gain and damping, flies compensated for changes in inertia with only minor changes in performance at the cost of overall stability and a larger response time. Such compensatory changes could not be explained by feedback alone (Figure 2A), nor could they be achieved using feedback from one sensory modality (Figure 3 & 4). Flies adjusted the initial torque to generate compensatory saccades to added inertia, suggesting that they modulate internal control commands and sense the extra mechanical load. We propose a control scheme which is composed of two feedback loops: a nested loop is driven by mechanosensory feedback that regulates yaw damping and an outer loop that regulates visuomotor gain (Figure 4). Taken together, our findings indicate that flies modulate neural controller gains to maintain performance at the expense of increased response time.

### Flies compensate for added inertia by trading-off response time

Magnetically tethered flies with added inertia suffered only a marginal drop in gain, but a significant drop in phase during gaze stabilization. When the yaw inertia of flies was increased by 32X, flies began to exhibit a peak in closed-loop (behavioral) gain which is characteristic of underdamped second order systems, indicating that performance had begun to suffer as the gain was larger than unity around this peak (Figure 2B). The underlying reason behind this shift in dynamics is not intuitive. The dip in phase became more prominent at higher inertia and was a direct result of changes in the system time delay. Indeed, simulations indicated that increasing the time delay alone captured the observed changes in the frequency response of flies following addition of inertia (Figure 2C). At present the underlying mechanism driving this change in time delay remains obscure. Changes in time delay could be a manifestation of some higher order dynamics that cannot be modeled using the current framework. Work that investigated flower tracking in hawkmoths in environments with different levels of luminance found that lower levels of light resulted in larger time delays, which can be modeled by a change in the low-pass filter time constant of visual processing (Sponberg et al., 2015). While the body dynamics of moths were not modified in that study, this study hints that the observed changes in delay in fruit flies may be due to active neural modulation. Alternatively, the increase in time delay could be a result of an increase in reaction time, that is a consequence of the fly compensating for an unusual perturbation. Indeed, larger time delays can negatively impact system yaw stability in insect flight (Elzinga et al., 2012). Taken together, in response to added inertia, flies maintain roughly the same gaze stabilization performance at the expense of stability and response time.

Our paradigm allowed us to push the performance of flies beyond natural conditions as lift generation and yaw stabilization were decoupled, which here we used to reveal the capacity of the nervous system to adapt yaw steering. Indeed, in free flight, flies with 1X or 2X added inertia may be more naturalistic. Nevertheless, the ability of flies with large added inertia to stabilize gaze in the magnetic tether is a strong indication of the capacity for adaptive compensatory behavior. Our results should be interpreted with appropriate caution as the tethering paradigm restricts the motion of flies to rotation about the yaw axis. This is unnatural for flies, although they can perform nearly pure yaw rotation in free flight (Bergou et al., 2010). Further, the tether supports the weight of the fly and cylinder which eliminates the need for lift generation. As a result, the wing kinematics of magnetically tethered flies likely deviate from those in free flight.

### Flies adaptively control saccade dynamics

Increasing the yaw inertia of flies altered saccade dynamics (Figure 5). By modeling the yaw dynamics of tethered flies as a first order system, we could predict how saccade dynamics should change in the absence of sensory feedback and yaw torque modulation (Figure 5A). By assuming flies produced the same yaw torque regardless of inertia treatment, the model predicted that saccades should exhibit drastically smaller displacements, peak velocities, and durations (Figure 5A lower panel). This is in stark contrast to empirical data (Figure 5A upper panel). In fact, the average velocity of saccades at different added inertias did not remotely resemble our predicted results (Figure 5A). Thus, differences between model and data suggest a mechanism that modulates saccade dynamics due to mechanical loading. Previous work found that altering haltere feedback had a significant impact on saccade dynamics (Bender and Dickinson, 2006b). Therefore, one possibility is that changes in saccade dynamics are a result of changes in haltere gyroscopic feedback due to alterations in body inertia. This hypothesis also presumes that the fly has some internal model or, alternatively, a goal at the start of the saccade and relies on mechanosensory feedback to achieve this goal. Work measuring the torque production of rigidly tethered flies during a saccade reported durations as high as 500 ms (Tammero and Dickinson, 2002); much larger than anything reported in freely flying or magnetically tethered flies (Cellini and Mongeau, 2020b). This also suggests that contrary to previous findings, flies employ mechanosensory feedback not only when ‘braking’ during a saccade, but also to modulate saccade initiation. Alternatively, flies may have updated an internal model which accounted for the added inertia. By comparing the saccade torque profile of flies with added inertia to unaltered flies, we found a clear increase in torque production with increasing inertia (Figure 5B). While much larger in magnitude, the torque profile of flies with added inertia resembled that of intact flies, which suggests that changes in saccade dynamics may be a result of mechanosensory feedback instead of learning. Further supporting this conclusion is the observed elevation in saccade duration. Intriguingly, humans similarly compensate for artificially increased inertia during rapid rotational maneuvers (Lee et al., 2001).

### Flies maintain stability by combining sensory feedback with adaptive control

By combining experiments with simulation, we found that increasing inertia had little impact on gaze stabilization performance. While subject to some changes, the open-loop gain and pole locations did not deviate as much as predicted from simulation (Figure 3D,E). Similarly, the estimated open-loop gain and time constant did not considerably vary with added inertias (Figure 3E,F). Such a feat was accomplished by increasing the effective yaw damping and visuomotor gain (Figure 3B,C). Using simulations, we found that damping must be actively modulated using neural control as passive damping alone cannot produce enough damping (Figure 4).

Based on this finding, we propose an adaptive control strategy which allows flies to regulate damping and visuomotor gain using multiple feedback loops. As flies integrate visual and mechanosensory feedback to stabilize flight (Sherman and Dickinson, 2004), we posit that flies regulate damping using a nested loop driven by mechanosensory feedback, whereas gain is regulated through an outer loop using vision (Figure 4A) (Elzinga et al., 2012). Using this scheme, flies can regulate damping by changing the haltere gain, whereas the open-loop gain regulates the visuomotor performance of the system. Had flies only regulated yaw damping in response to an increase in inertia, the optomotor response would have suffered a significant decrease in gaze stabilization gain at all frequencies (Figure S7). This is in stark contrast to our experimental results. Hence, tuning the visuomotor gain enabled flies to reduce the overall impact of added inertia by regulating the amount of torque produced. Through simulations, changes in inertia predicted an overall decrease in gain, even in the presence of feedback and in the absence of any modulation in internal gains (Figure 2A), thus we can conclude that flies likely implement an adaptive control scheme to compensate for changes in inertia. However, how flies regulate these internal parameters is not clear. We speculate that flies may implement an adaptive control scheme similar to a Model Reference Adaptive Scheme (MRAS) (Åström and Wittenmark, 2008). In this scheme, a system regulates the input to the plant (e.g., fly body) by comparing the observed output to reference output generated from a desired model. This hypothesis does not rule out that flies implement another adaptive scheme, or even a parallel robust scheme that relies solely on feedback. It is also possible that flies rely solely on a robust control scheme that contains a number of nested feedback loops which cannot be modeled by our current framework. Alternatively, flies may have learned a new controller altogether. In *de novo* learning, it is possible to change the entire controller to map sensory input to motor output (Yang et al., 2021). Overall, our results hint that flies implement an adaptive control scheme regulated by nested feedback loops to mitigate changes in inertia.

## Materials and Methods

### Animal preparation

Animal preparation was previously described in another study (Salem et al., 2020). Briefly, female fruit flies *Drosophila melanogaster* aged 3–5 days were cold anesthetized at 4°C using a Peltier cooling stage. Flies were then glued to a pin under a microscope and left to rest for approximately one hour before the start of experiments. The yaw inertia was altered by gluing a 3D printed cylinder onto the stainless-steel pin (Figure 1B). After the rest period, flies were suspended between two magnets and placed inside a virtual reality arena (Figure 1A) (Reiser and Dickinson, 2008). This configuration enables tethered flies to rotate about the yaw axis while restricting motion in the other directions. The pin’s yaw inertia was less than 1% that of the fly’s inertia. Hence, the pin did not introduce any significant inertia that may alter the interpretation of the collected data (rod diameter = 100 μm, tip diameter = 12.5 μm; Minutien pin, Fine Science Tools), as previously demonstrated (Cellini et al., 2022). Only flies that successfully completed at least three trials were used in subsequent analysis. Flies that continuously stopped flying during experiments or had very low baseline wingbeat amplitude (less than 100°) were not used in the analysis.

### Cylinder design and printing

Seven cylinders were designed to have progressively larger inertias that were integer multiples of the yaw inertia of fruit flies (5.2×10^-13^ kg m^2^ (Bender and Dickinson, 2006a)). To sample across a wide range of inertias and push the limit of flight performance, the cylinders were designed with logarithmically increasing yaw inertia (Table S5). The smallest cylinder had approximately the inertia of a single fly, whereas the inertia of the largest cylinder was around sixty-four times that of a fly. To ensure the inertia of cylinders closely matched the desired value, we 3D printed the cylinders using a resin 3D printer with a tolerance of 25 μm (Formlabs Form 3+ SLA printer). The cylinders were printed using a clear resin that had a density of 1.12–1.15 g/cm^3^. Due to the limitation in printer resolution, the actual inertias of the printed cylinders were slightly larger than designed. However, the actual mass and inertia of all cylinders fell within 10% of the desired inertia. Larger inertias (128X) were printed but not used in this study as the magnetic tether system could no longer support the extra weight of these cylinders. To ensure the cylinders were not a significant source of damping due to air friction, we estimated the torque due to air friction at different angular velocities (See Supplementary Material). Our calculations indicate the torque due to air friction is roughly two orders of magnitude smaller than the torque required to overcome yaw damping (Figure S8), thus providing assurance that air friction of the cylinders was not a significant source of damping.

### Stimuli and experimental setup

Using a virtual reality arena, we presented magnetically tethered flies with a visual stimulus (moving background) that elicited an optomotor response. The background consisted of uniformly spaced bars with a spatial wavelength of 22.5° subtending onto the fly eye. To test the impact of increasing yaw inertia on flight performance, we presented magnetically tethered flies with a visual sum-of-sines stimulus (Figure 1). This stimulus was generated by adding nine sine signals with distinct frequencies that ranged from 0.35 Hz to 13.7 Hz. Each component of this stimulus had a random phase and an amplitude normalized to a velocity of 52° s^-1^. This ensured the stimulus velocity did not saturate the visual and motor systems, as previously described (Cellini et al., 2022). Each trial lasted 20 seconds and was presented five times to each fly. A second set of experiments was conducted to measure the impact of increasing inertia on the yaw stability of flies in the presence of a statis stimulus. Flies were presented with the same uniform background which was kept stationary for 10 second and underwent five trials. Flies that did not complete more than three trials or had a low wingbeat amplitude (less than 100°) were not used in the analysis. Changes in heading of the flies were measured using a bottom view camera (Basler acA640–750um) recording at 80–100 frames per second (fps). Wing data was collected by measuring the wingbeat amplitude (extreme position at downstroke-to-upstroke reversal) using a modified version of Kinefly (Suver et al., 2016). To enable accurate measurements of wingbeat amplitude, the bottom view videos were registered with respect to the fly’s reference frame prior to tracking the wings.

### Tracking in the magnetic tether

The head and body motion were tracked using a custom MATLAB code that has been previously described (Cellini et al., 2022). The amplitudes of both wings were estimated by measuring the angle the edge of the wing blur made with the axis of the fly’s body. Estimates of the wingbeat amplitude were measured using a modifies version of Kenifly (Suver et al., 2016). Prior to measurement of wing and head kinematics, videos were registered to eliminate the yaw motion of the body, as done previously (Cellini et al., 2022).

### Flight performance metric

The impact of adding inertia was measured using multiple performance metrics commonly used in the system identification of engineering systems. The system identification analysis was conducted using MATLAB, and each metric was estimated for individual flies and then averaged out across all flies to determine the grand mean for each inertia treatment. The gain was calculated by dividing the FFT magnitude of the fly’s heading (output) with that of the visual stimulus (input). The phase difference was estimated by subtracting the output’s phase from that of the input. The coherence was estimated using the MATLAB built-in function *mscohere*. Finally, we used the compensation error as an overall metric to measure changes in performance (Roth et al., 2011). The compensation error is a metric that combines gain and phase to indicate how well flies compensate for a moving background. A gain of unity and a phase difference of zero produce zero compensation error and indicate perfect tracking. The compensation error is calculated by finding the vector distance in the complex plane (norm) between the actual tracking performance *H* and the perfect tracking *Z*_0_ and can be expressed as

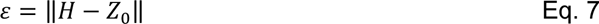

Therefore, a compensation error of 0 indicates that flies perfectly compensated for the visual stimulus, a compensation error between zero and one indicate imperfect compensation, and values greater than one indicate that the system can a better job at stabilizing the input by effectively not responding. To avoid phase wrapping, the averaged phase difference was calculated using the circular statistics toolbox in MATLAB (Berens, 2009).

### Transfer function fitting and system identification

The magnetic tether restricts the body motion of fruit flies to rotation about the axis of the pin (yaw axis). Therefore, we can approximate the yaw dynamics using the following equation

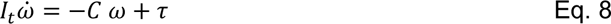

where *I_t_* is the total yaw inertia of the fly (body inertia plus cylinder inertia), *C* is the yaw damping coefficient, τ is the torque generated by the wings, and ω is yaw angular velocity of the fly (Cellini et al., 2022; Salem et al., 2022). Transfer function fitting was performed using a custom designed MATLAB code and the method is detailed elsewhere (Roth et al., 2012). In short, we fit the error/output data to a first order transfer function in Equation 2.

We did not fit the value of the inertia, rather we assumed a constant value for each group (total inertia = fly inertia + inertia of cylinder). These parameters were estimated using a least square estimate as described in a previous study (Roth et al., 2012). Only fits with a at least 65% goodness of fit (GoF) were used in the transfer function fitting and parameter estimation. The GoF was at least 84 % for all groups and detailed estimate for each inertia treatment is in Table S6 and Figure S4B. The FRF obtained from flies with an added inertia of sixty-four times was not used in transfer function fitting due to the low overall coherence of the response.

### Flapping-counter torque estimates

In this study, we found that flies modulated yaw damping in response to changes in inertia to maintain performance. However, it was not clear if damping is actively modulated using neural control, or a byproduct of changes in wing kinematics aimed at elevating torque production. To determine the nature of this change, we estimated the flapping counter-torque (FCT), which is a passively generated torque in flapping flight that is produced during turns (Cheng et al., 2010). In the magnetic tether, the FCT counter-acts rotations about the yaw axis, thus, it can be thought of as viscous damping about the yaw axis proportional to yaw angular velocity. The method for estimating FCT has been described previously (Salem et al., 2022). Briefly, we estimated the stroke angle of flies by multiplying the base stroke angle from free flight data (Muijres et al., 2017a) with a correction factor, which was then projected onto the stroke plane. For rotation angles, we used the intact baseline rotation angles measured in free flight (Muijres et al., 2017a). A comprehensive derivation of the FCT equations can be found in (Cheng et al., 2010). The wing morphological parameters required to calculate the FCT were estimated using images of wings taken under a microscope and analyzed using a custom MATLAB code. To estimate how changes in wing kinematics altered the FCT, we estimated the FCT for a flapping frequency ranging from 200 Hz to 1000 Hz. We also modified the rotation angle by multiplying the baseline rotation angle for both wings with a scaling factor. The passive damping was then estimated for different combinations of flapping frequency and rotation angles (Figure 4D). This allowed us to determine if changes in wing kinematics could produce a large enough yaw damping from the FCT model alone.

### Saccade detection and analysis

In this study, we only estimated the dynamics of spontaneous saccades. Unlike reset and catch-up saccades which are triggered by an external visual stimulus (Cellini and Mongeau, 2020b; Cellini et al., 2021; Mongeau and Frye, 2017; Mronz and Lehmann, 2008), spontaneous saccades are internally triggered (Censi et al., 2013). Hence, we only used saccade data generated from our static background experiments. Saccade detection was accomplished by using methods previously described (Mongeau and Frye, 2017). Magnetically tethered flies began to oscillate about the pin’s axis when yaw inertia was increased by more than 8X, which complicated automatic saccade detection as the dynamics of the oscillations were close to the dynamics of saccades. To ensure no false saccades were included, we designed custom code (MATLAB) which flagged saccades with a displacement smaller than 10° and with a duration smaller than 50 ms. We manually verified and removed flagged saccades to confirm their identity via a custom graphical user interface. Further complicating the comparison of saccade dynamics was the large sample size (>100 saccades per group), thus a tiny difference in saccade dynamics produces small *p* values. Therefore, a comparison may yield a statistically significant, but not a biologically relevant difference. To address this issue, we computed Hedge’s g, which presents a metric of effect size independent of sample size (Hedges, 1981). This allowed us to properly compare changes in saccade dynamics of the inertia added flies to that of the unaltered group.

### Statistics and comparison

For all box plots, the central line is the median, the bottom and top edges of the box are the 25th and 75th percentiles and the whiskers and extend to ±2.7 standard deviations. Unless otherwise specified, we report means ± 1 standard deviation. Significant differences are stated as \**p* ≤ 0.05, \*\**p* ≤ 0.01, \*\*\**p* ≤ 0.001. Unless otherwise noted, saccade dynamics were compared using the effect-size model Hedge’s g.

## Author Contributions

W.S. and J.M.M. designed research; W.S. and E.J. performed research; B.C. contributed new analytic tools; W.S. and B.C. analyzed data; and W.S. and J.M.M. wrote the paper.

## Data availability statement

All code and data will be made available on Penn State ScholarSphere.

## Supplementary Material

### Estimate of rotating cylinder torque

To estimate the friction torque on a rotating cylinder, we assumed a solid cylinder rotating at a constant velocity. We estimated the resulting friction torque for the different cylinder inertias with distinct geometry (Table S5). For each cylinder, we calculated the Reynolds number (*Re*) for angular velocities ω ranging from 10–300 °s^-1^ with a kinematic viscosity of air of 1.57×10^-5^ m^2^/s, which yielded a flow in a laminar regime. For a rotating cylinder in a laminar flow [1], the moment coefficient is given by

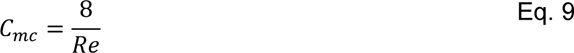

Thus, the torque to overcome friction drag for a rotating cylinder is

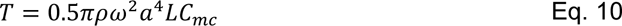

where *ρ* is the air density (1.225 kg/m^3^), ω is the angular velocity, *a* is the radius, and *L* is the length of the cylinder.

**Figure S1.**
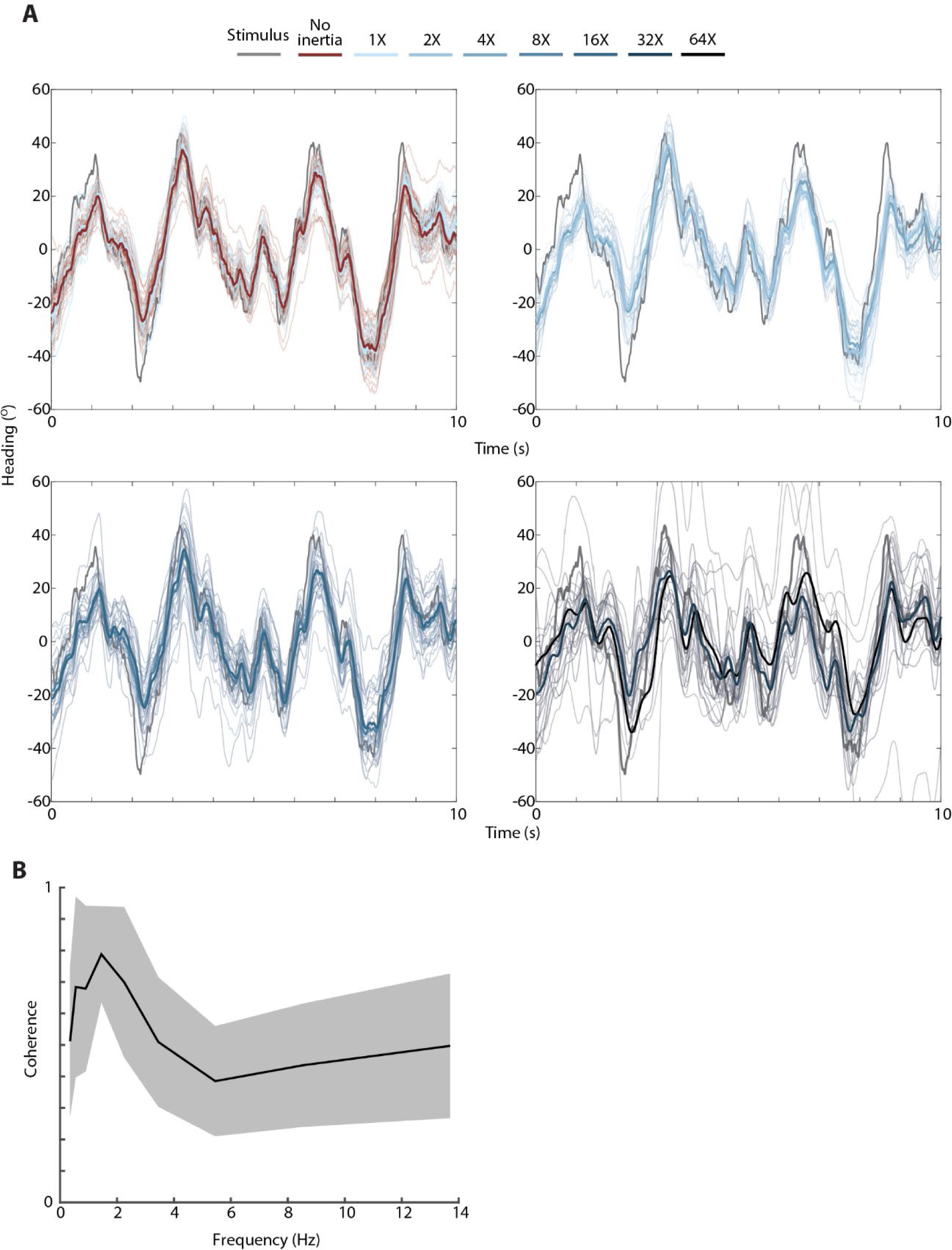
Individual and average response of flies with different amount of added inertia. A) Individual (light lines) and averaged response (solid lines) to a sum-of-sines stimulus. B) The average coherence of flies with 64X added inertia. No inertia added: *n* = 41 flies; 1X: *n* = 11 flies; 2X: *n* = 15 flies; 4X: *n* = 19 flies; 8X: *n* = 17 flies; 16X: *n* = 17 flies; 32X: *n* = 8 flies. Shaded area: ± 1 STD.

**Figure S2.**
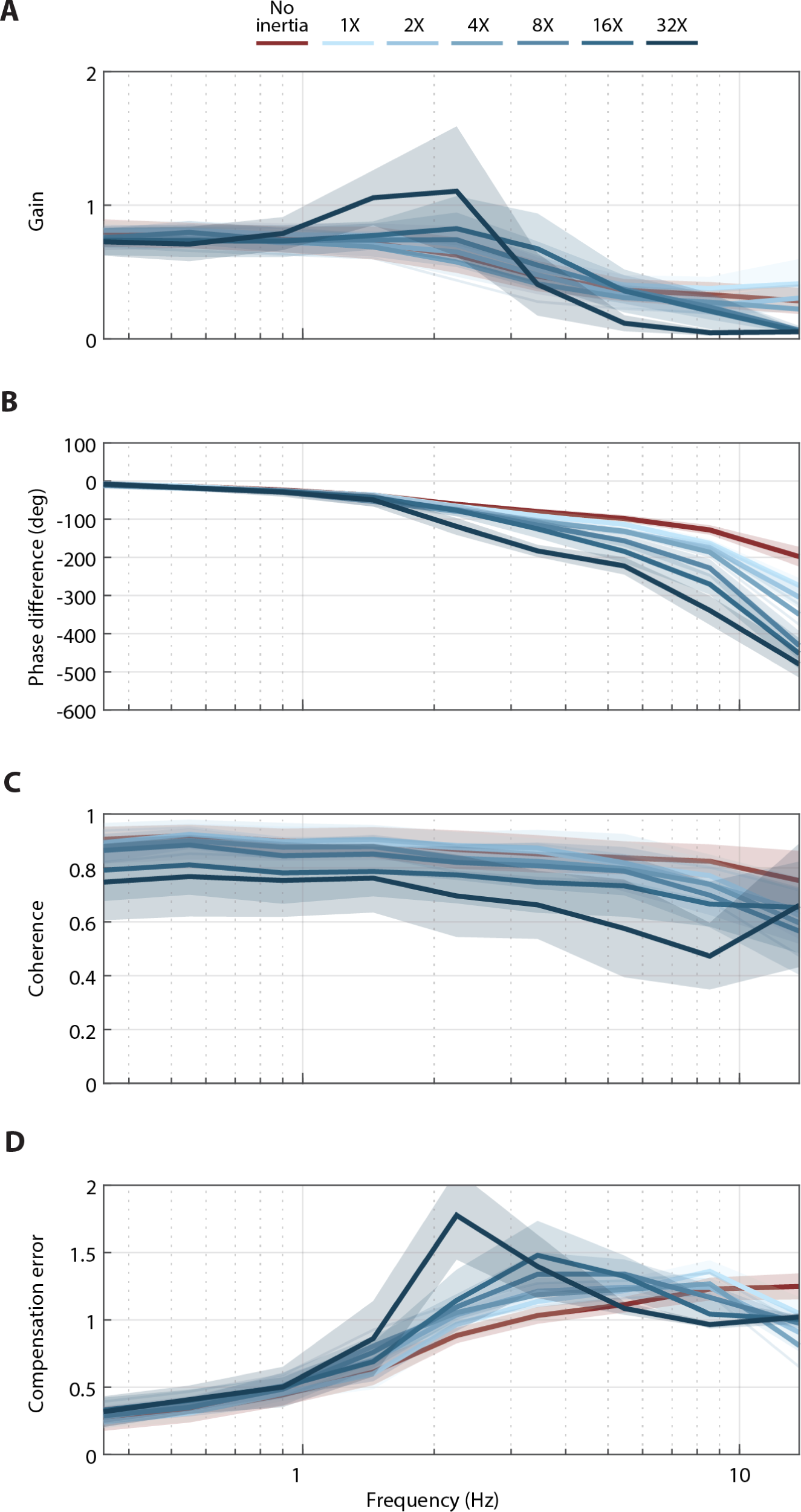
The average frequency domain response (solid lines) of flies with different amount of added inertia. A) Gain. B) Phase difference. C) Coherence. D) Compensation error. No inertia added: *n* = 41 flies; 1X: *n* = 11 flies; 2X: *n* = 15 flies; 4X: *n* = 19 flies; 8X: *n* = 17 flies; 16X: *n* = 17 flies; 32X: *n* = 8 flies. Shaded area: ±1 STD.

**Figure S3.**
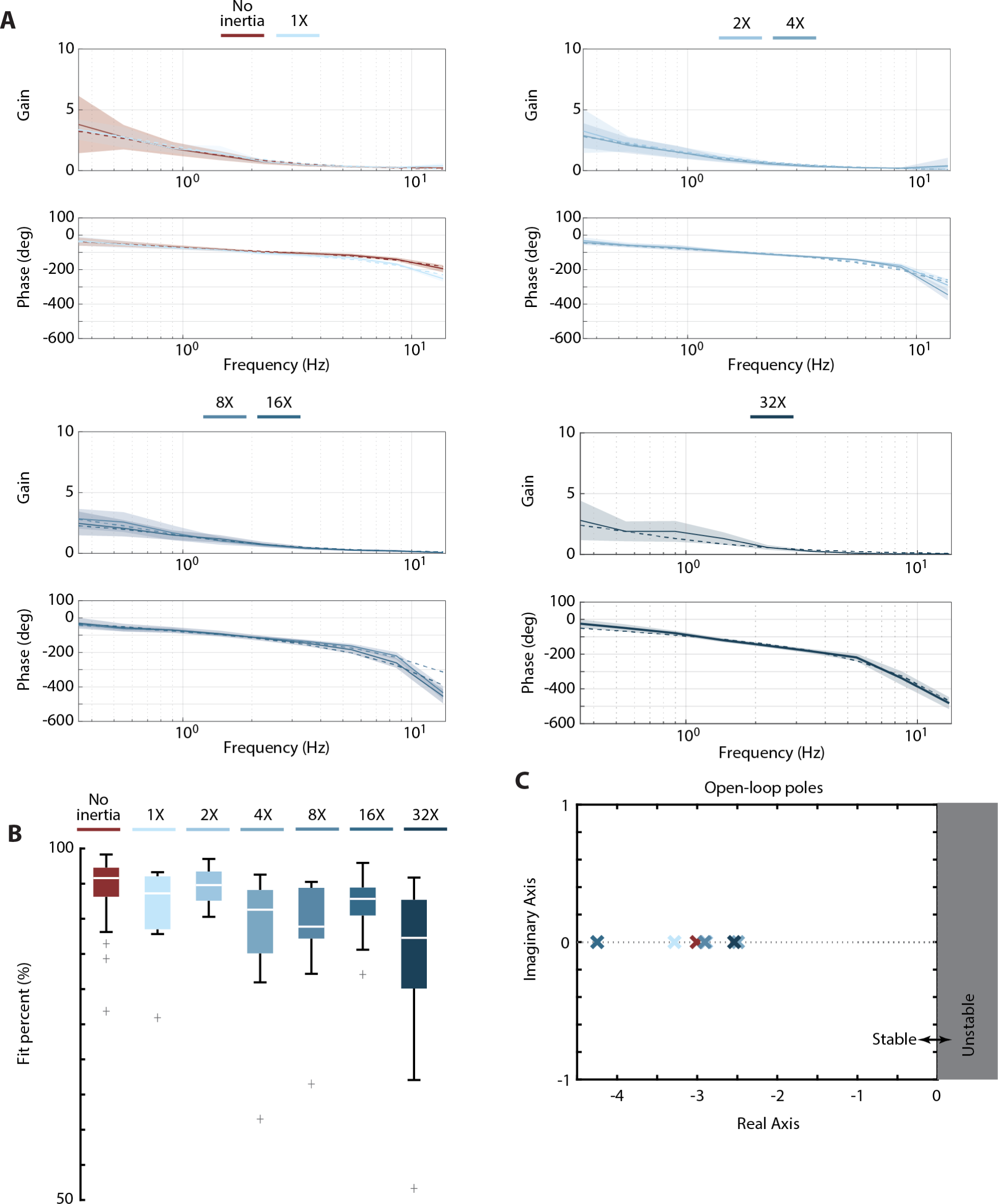
Open-loop Bode plots and best-fit transfer functions to experimental data. A) Open-loop Bode plots (solid lines) with the best-fit transfer function (dashed line). Shaded area: ±1 STD. For visual clarify, two sets of inertia at most are shown on each plot. B) Fit percentage of each data set to the first-order transfer function ***G*(*s*)** (Eq. 2). C) Average pole location of the open-loop transfer functions. No inertia added: *n* = 41 flies; 1X: *n* = 11 flies; 2X: *n* = 15 flies; 4X: *n* = 19 flies; 8X: *n* = 17 flies; 16X: *n* = 17 flies; 32X: *n* = 8 flies.

**Figure S4.**
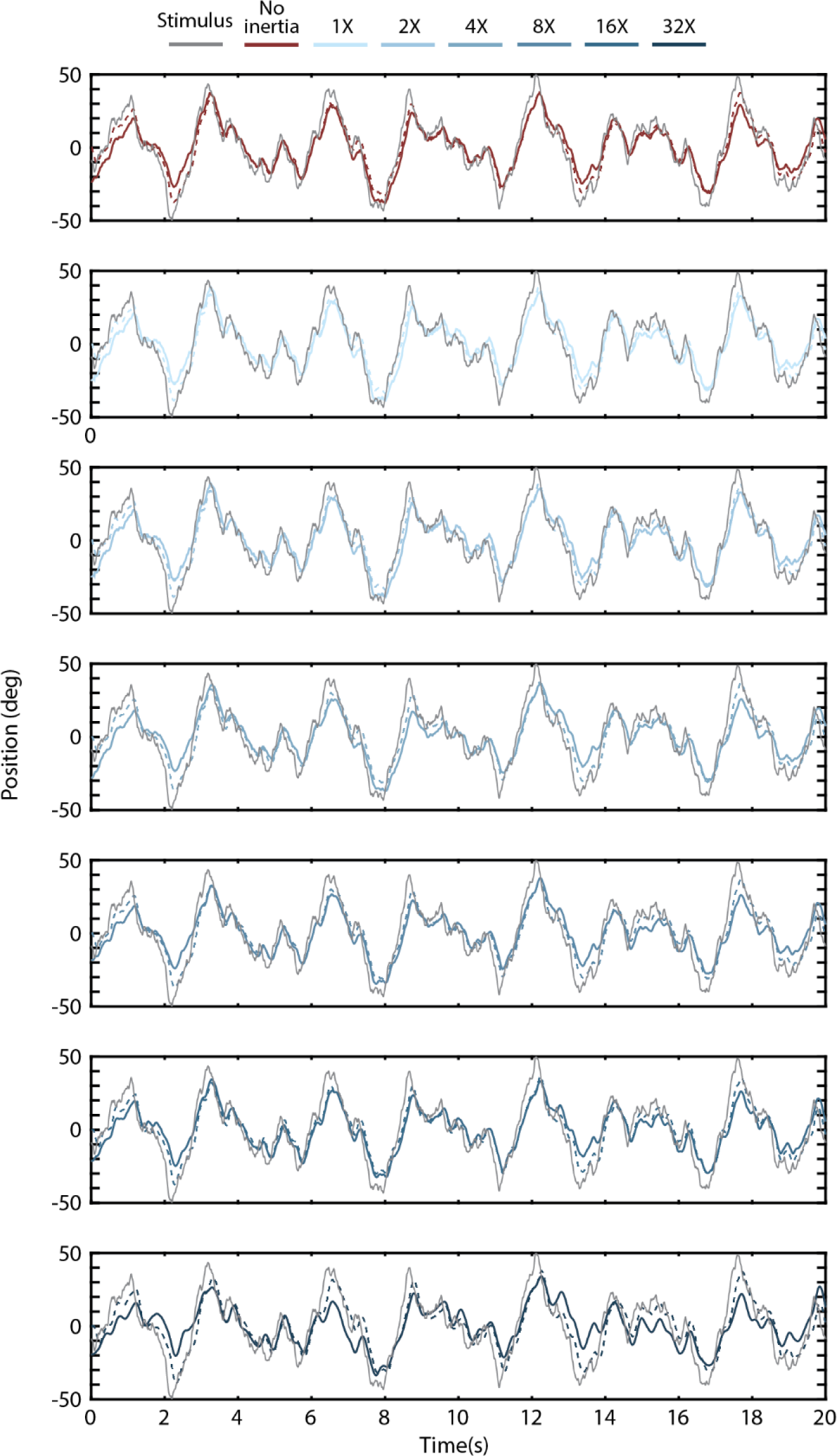
Simulated (dashed lines) and actual mean response (solid lines) of inertia altered flies to the sum-of-sines visual stimulus (grey). No inertia added: *n* = 41 flies; 1X: *n* = 11 flies; 2X: *n* = 15 flies; 4X: *n* = 19 flies; 8X: *n* = 17 flies; 16X: *n* = 17 flies; 32X: *n* = 8 flies.

**Figure S5.**
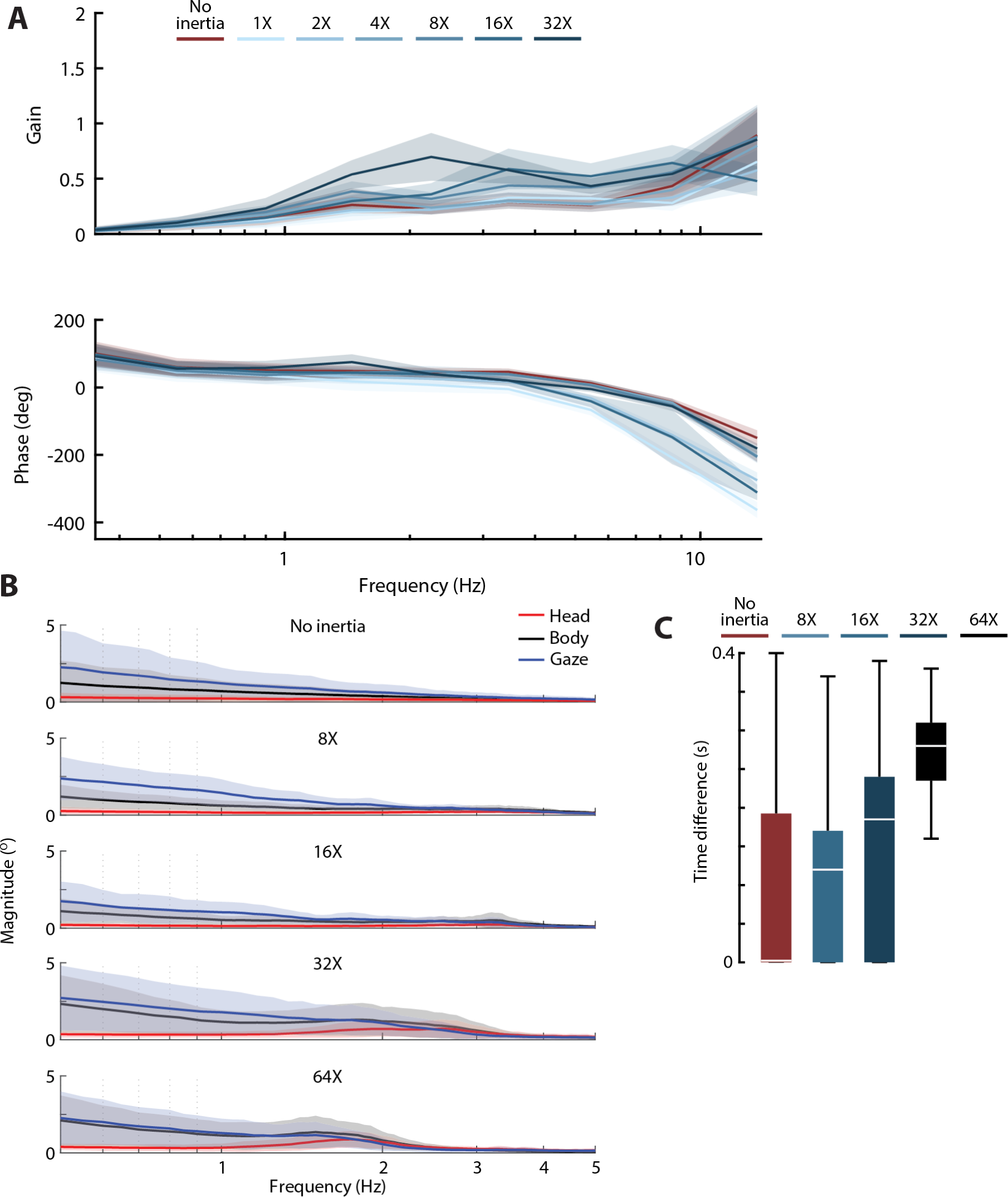
Impact of increasing body inertia on head motion in the presence and absence of a moving stimulus A) The frequency response of the head for a visual sum-of-sines stimulus. B) A magnitude plot of head oscillations for magnetically tethered flies presented with a static stimulus. C) Time difference between the head and body response to a static background. No added inertia: *n* = 13 flies; 16X inertia: *n* = 7 flies; 32X inertia: *n* = 9 flies; 64X inertia: *n* = 13 flies.

**Figure S6.**
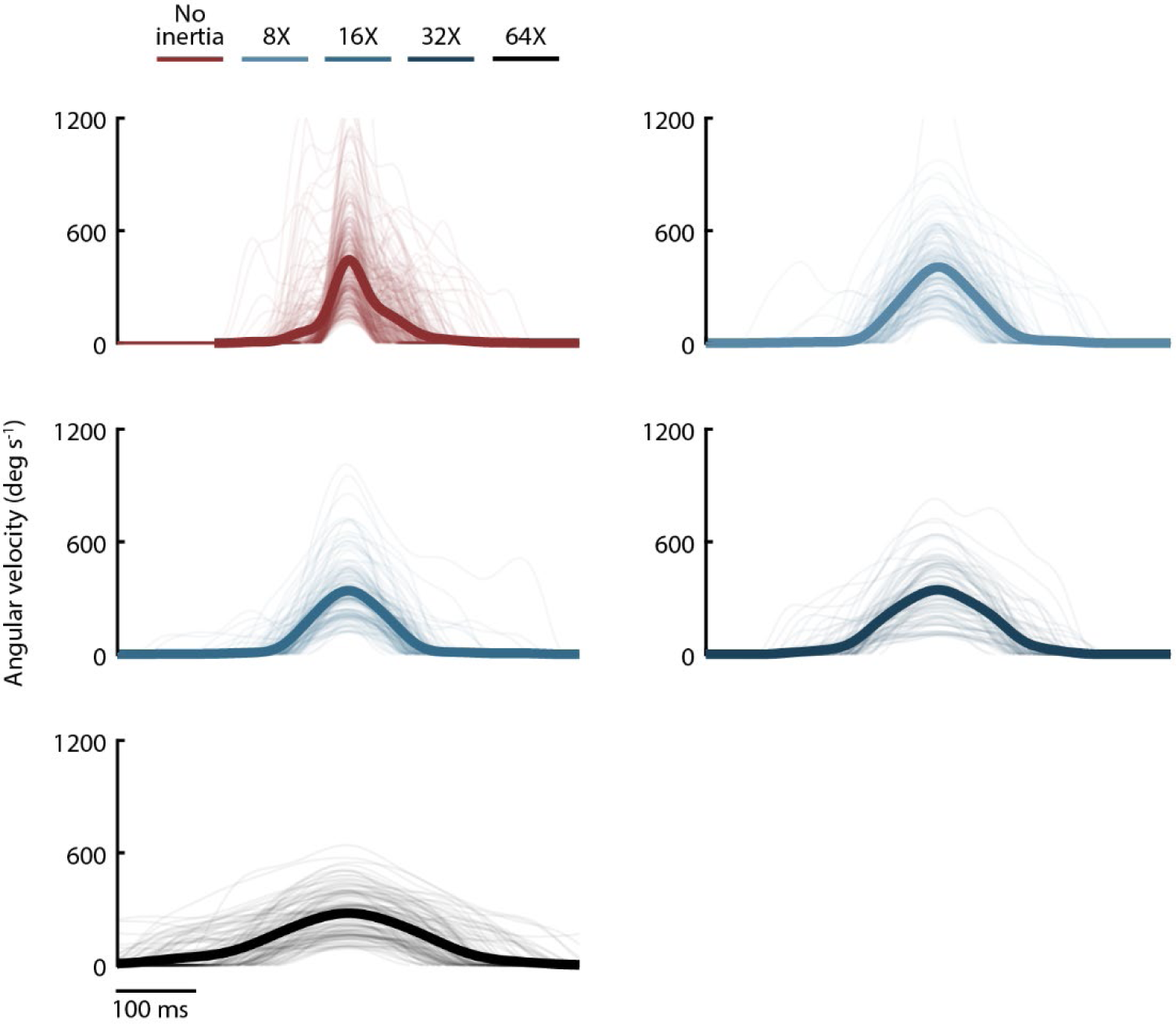
Average saccade velocity profiles (solid lines) along with individual saccades (thin lines). No added inertia: *n* = 301 saccades from 13 flies; 8X: *n* = 148 saccades from 9 flies; 16X: *n* = 139 saccades from 7 flies; 32X: *n* = 89 saccades from 9 flies; 64X: *n* = 118 saccades from 13 flies.

**Figure S7.**
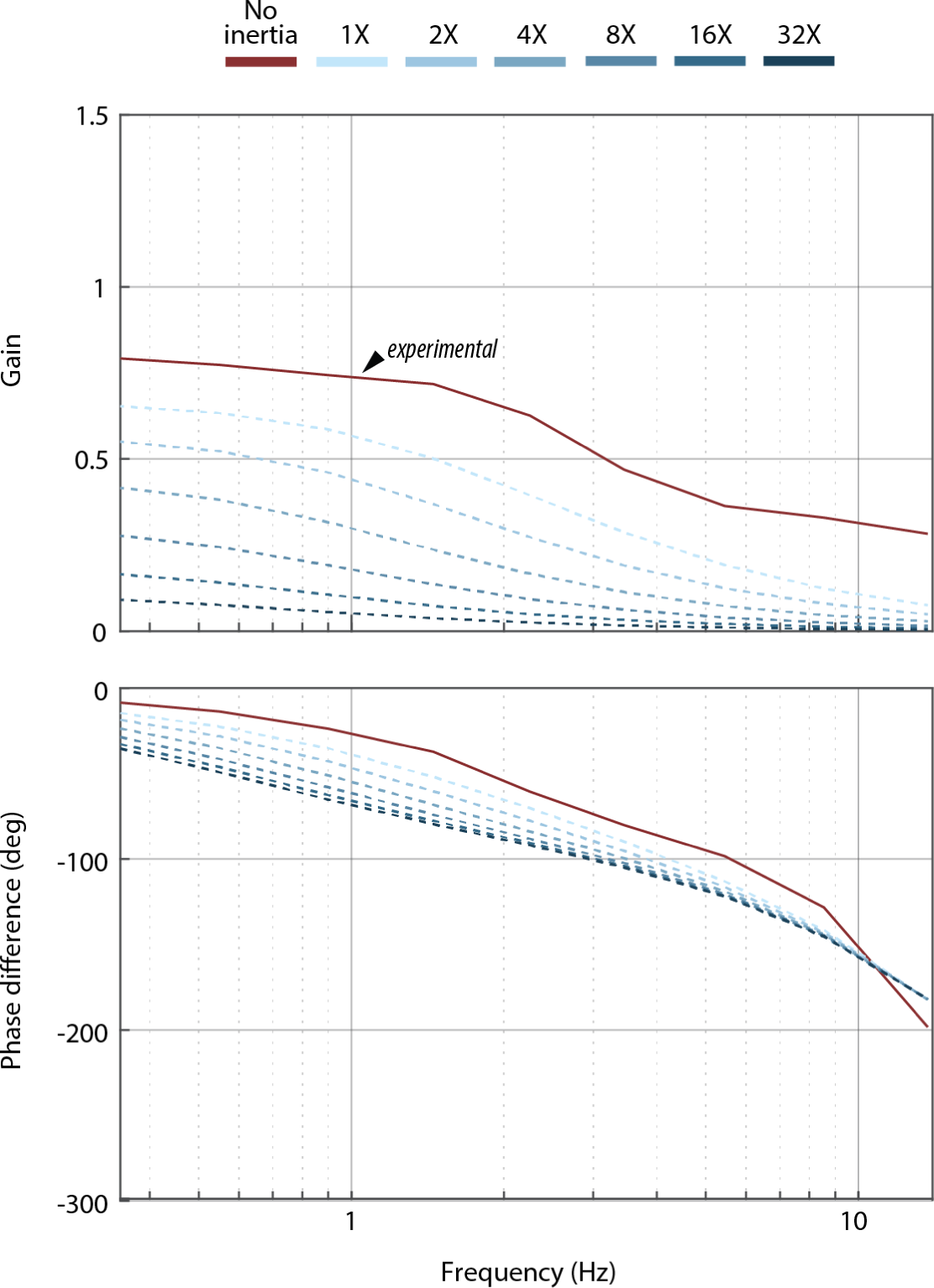
Predicted closed-loop frequency response functions if flies only altered damping in response to changes in inertia. Prediction: dashed lines. Empirical frequency response function for no-added-inertia flies: solid line. The simulation predicts a drastic drop in gain at all frequencies.

**Figure S8.**
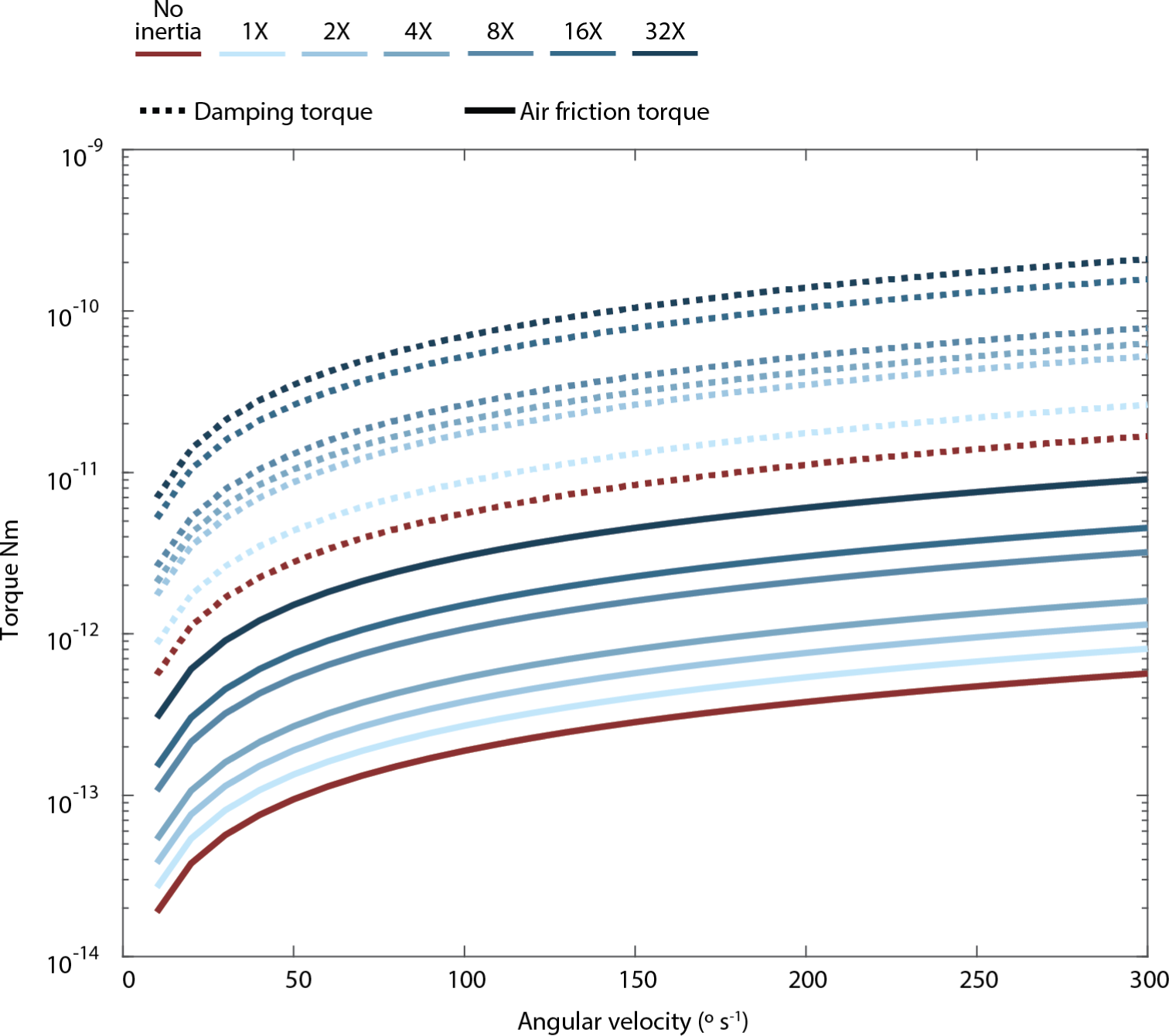
Aerodynamic torque (solid line) and the resulting flapping counter-torque (passive damping; dashed lines) that are produced due to rotation about the yaw axis. The yaw damping of the fly is approximately two orders of magnitude larger than the friction torque produced by the cylinder rotation.

**Table S1.** Gain (average). p values computed from ANOVA, DOF = 6. No inertia added n = 41 flies; 1X n = 11 flies; 2X n = 15 flies; 4X n = 19 flies; 8X n = 17 flies; 16X n = 17 flies; 32X n = 8 flies.

| Frequency | 0.35 | 0.55 | 0.9 | 1.45 | 2.25 | 3.45 | 5.45 | 8.55 | 13.7 |
| --- | --- | --- | --- | --- | --- | --- | --- | --- | --- |
| Inertia |  |  |  |  |  |  |  |  |  |
| 0 | 0.792 | 0.773 | 0.744 | 0.718 | 0.626 | 0.469 | 0.364 | 0.33 | 0.283 |
| 1 | 0.812 | 0.813 | 0.781 | 0.773 | 0.704 | 0.52 | 0.403 | 0.377 | 0.412 |
| 2 | 0.760 | 0.761 | 0.721 | 0.704 | 0.65 | 0.479 | 0.321 | 0.248 | 0.304 |
| 4 | 0.759 | 0.746 | 0.731 | 0.688 | 0.574 | 0.412 | 0.307 | 0.274 | 0.223 |
| 8 | 0.756 | 0.797 | 0.738 | 0.739 | 0.741 | 0.551 | 0.369 | 0.238 | 0.064 |
| 16 | 0.737 | 0.734 | 0.731 | 0.773 | 0.825 | 0.674 | 0.358 | 0.207 | 0.055 |
| 32 | 0.726 | 0.709 | 0.788 | 1.056 | 1.105 | 0.406 | 0.116 | 0.045 | 0.051 |
| **p-value** | **0.122** | **0.081** | **0.48** | **1.7e-14** | **2.6e-9** | **1.2e-4** | **1.1e-13** | **1.6e-18** | **1.6e-23** |

**Table S2.** Phase (average). p values computed from ANOVA, DOF = 6. No inertia added n = 41 flies; 1X n = 11 flies; 2X n = 15 flies; 4X n = 19 flies; 8X n = 17 flies; 16X n = 17 flies; 32X n = 8 flies.

| Frequency | 0.35 | 0.55 | 0.9 | 1.45 | 2.25 | 3.45 | 5.45 | 8.55 | 13.7 |
| --- | --- | --- | --- | --- | --- | --- | --- | --- | --- |
| Inertia |  |  |  |  |  |  |  |  |  |
| 0 | -8.46 | -13.65 | -23.69 | -37.24 | -60.58 | -80.11 | -98.37 | -128.3 | -198.95 |
| 1 | -6.7 | -13.68 | -25.33 | -36.57 | -65.77 | -91 | -113.88 | -163.19 | -273.93 |
| 2 | -7.48 | -17.36 | -28.21 | -44.9 | -74.08 | -103.98 | -132.94 | -174.52 | -304.36 |
| 4 | -11.84 | -19.19 | -28.56 | -53.35 | -79.39 | -106.56 | -131.78 | -186.68 | -349.92 |
| 8 | -9.27 | -17.71 | -27.23 | -49.53 | -75.64 | -115.84 | -156.77 | -227.61 | -434.17 |
| 16 | -8.44 | -17.96 | -27.34 | -42.74 | -76.74 | -125.97 | -184.58 | -269.82 | -450.38 |
| 32 | -8.32 | -17.94 | -29.54 | -49.48 | -118.93 | -183.38 | -222.75 | -340.02 | -478.95 |
| **p-value** | **0.53** | **0.0025** | **0.073** | **1.2e-6** | **6.1e-32** | **3.2e-52** | **4.1e-54** | **6.5e-67** | **3.4e-68** |

**Table S3.** Coherence (average). *p* values computed from ANOVA, DOF = 6. No inertia added *n* = 41 flies; 1X *n* = 11 flies; 2X *n* = 15 flies; 4X *n* = 19 flies; 8X *n* = 17 flies; 16X *n* = 17 flies; 32X *n* = 8 flies.

| Frequency | 0.35 | 0.55 | 0.9 | 1.45 | 2.25 | 3.45 | 5.45 | 8.55 | 13.7 |
| --- | --- | --- | --- | --- | --- | --- | --- | --- | --- |
| Inertia |  |  |  |  |  |  |  |  |  |
| 0 | 0.91 | 0.92 | 0.9 | 0.9 | 0.87 | 0.85 | 0.84 | 0.83 | 0.75 |
| 1 | 0.88 | 0.9 | 0.89 | 0.89 | 0.87 | 0.86 | 0.81 | 0.77 | 0.62 |
| 2 | 0.89 | 0.92 | 0.9 | 0.91 | 0.88 | 0.87 | 0.82 | 0.74 | 0.64 |
| 4 | 0.88 | 0.9 | 0.88 | 0.88 | 0.84 | 0.82 | 0.8 | 0.74 | 0.6 |
| 8 | 0.86 | 0.88 | 0.85 | 0.85 | 0.82 | 0.81 | 0.79 | 0.7 | 0.57 |
| 16 | 0.79 | 0.81 | 0.78 | 0.79 | 0.77 | 0.75 | 0.73 | 0.67 | 0.65 |
| 32 | 0.75 | 0.77 | 0.75 | 0.76 | 0.7 | 0.66 | 0.58 | 0.47 | 0.66 |
| **p-value** | **0** | **0** | **0** | **0** | **0** | **0** | **0** | **0** | **0** |

**Table S4.**
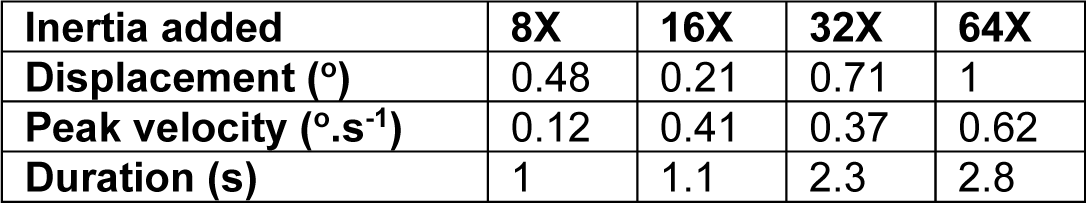
Hedge’s g for added inertia flies compared to the baseline no added inertia flies. No added inertia n = 301 saccades; 8X n = 148 saccades; 16X n = 139 saccades; 32X n = 89 saccades; 64X n = 118 saccades

**Table S5.** Cylinder designed and actual average inertias along with the cylinder sizes

| Inertia factor | Designed inertia (kg·m<sup>2</sup>) | Measured inertia (kg·m<sup>2</sup>) | Height (mm) | Outer Diameter (mm) | Inner Diameter (mm) |
| --- | --- | --- | --- | --- | --- |
| 1 | $5.2 \times 10^{-13}$ | $5.75 \times 10^{-13}$ | 0.70 | 1.60 | 0.60 |
| 2 | $10.4 \times 10^{-13}$ | $11.3 \times 10^{-13}$ | 0.70 | 1.91 | 0.60 |
| 4 | $20.8 \times 10^{-13}$ | $22.8 \times 10^{-13}$ | 0.70 | 2.27 | 0.60 |
| 8 | $41.6 \times 10^{-13}$ | $42.2 \times 10^{-13}$ | 0.70 | 2.69 | 0.60 |
| 16 | $83.2 \times 10^{-13}$ | $84.6 \times 10^{-13}$ | 1.40 | 2.69 | 0.60 |
| 32 | $166 \times 10^{-13}$ | $167 \times 10^{-13}$ | 1.40 | 3.20 | 0.60 |
| 64 | $332 \times 10^{-13}$ | $334 \times 10^{-13}$ | 2.80 | 3.20 | 0.60 |

**Table S6.** Percent goodness of fit for different inertias. Mean ± 1 STD.

| Inertia factor | Goodness of fit (GoF) % |
| --- | --- |
| 0 (no added inertia) | $94.5 \pm 4.27$ |
| 1 | $91.1 \pm 7.00$ |
| 2 | $94.6 \pm 2.34$ |
| 4 | $88.2 \pm 8.80$ |
| 8 | $88.8 \pm 6.73$ |
| 16 | $92.0 \pm 4.27$ |
| 32 | $84.0 \pm 2.34$ |
| 64 | Data not fit |

**Movie S1.** Oscillations for a magnetically tethered fly with 64X added inertia presented with a static stimulus. Top left: Bottom view of a single fly within animated virtual reality arena during a full 20 s trial. Top right: Same as top left but shown in a body reference frame. Bottom: stimulus (green) and body (magenta). The arena is not drawn to scale for visual clarity. The cylinder in the body fixed flew appears off-axis due to camera perspective errors, i.e., the lens is not perfectly perpendicular to the fly and added cylinder. The video was recorded at 100 fps but showed at 50 fps.

**Movie S2**. Same as Movie S1 but for a fly with 2X added inertia presented a sum-of-sines stimulus. The video was recorded at 100 fps but showed at 50 fps.

